# A case for multi-gear assessments: Detection probabilities of nearshore fish with eDNA and seine nets vary by functional traits

**DOI:** 10.1101/2025.11.28.691209

**Authors:** Lia K. Domke, Kimberly J. Ledger, Jessica M. Whitney, Shannon M. Kachel, Wes Larson, Ginny L. Eckert, Diana S. Baetscher

## Abstract

Ecological studies aim to understand species distributions, yet the methods used to sample affect which species are detected (i.e., gear selectivity) and may be influenced by species traits. Interactions between traditional gear selectivity for marine fishes and their species traits have been studied, but such studies have generally not extended to selectivity of environmental DNA (eDNA). Here, we investigated which functional species traits (scale type, schooling behaviour, and position-in-water-column) are selected for by eDNA metabarcoding and beach seines in nearshore eelgrass, mixed eelgrass, and understory kelp habitats. Using data from 35 sites across southeast Alaska, we applied occupancy modeling to estimate detection of species traits by each gear type. Detection probability with eDNA was 27 times greater for species with deciduous scales compared to species with non-deciduous scales, and lower for species with plates (rather than scales). Conversely, species with plates showed greater odds of detection with beach seines. Given the novelty of eDNA sampling, quantifying interactions between functional traits and gear selectivity will be important to accurately characterize species distributions across marine habitats.

## 1 Introduction

Ecological studies sample species communities to understand diversity, ecosystem function, and provide information about species distributions for management and conservation actions (Lefcheck et al., 2015; Margules and Pressey, 2000; Oliver et al., 2015). Yet the methods used to sample (i.e., gear type) can influence how effectively particular species are detected (i.e., selectivity; ICNAF, 1963). In aquatic ecosystems, nets have traditionally been used to sample the fish community, but comparisons between net sampling and alternative gear types show bias in the species of fishes captured by different gears (McClanahan and Mangi, 2004). More recently, environmental DNA (eDNA) has become a popular method for aquatic sampling, with many studies demonstrating increased species richness and diversity based on eDNA samples when compared to nets (He et al., 2022; Sard et al., 2019; Stoeckle et al., 2021). However, in most of these studies, eDNA does not detect all of the same fish species as the nets; thus, the suggestion is that complementary gear types should be used to maximize sampling efficiency and that eDNA should be one of the gear types employed in addition to traditional capture-based sampling methods (Andres et al., 2023; Gold et al., 2023; Stat et al., 2019).

Generally, species in eDNA samples are identified using metabarcoding, an efficient laboratory procedure that amplifies all DNA from a targeted taxonomic group (i.e., all teleost fishes). The amplification process results in species-specific bias originating from the unique characters in the DNA sequence for a given species and whether they perfectly match the primers used to bind to this sequence (Sato et al., 2021; Shelton et al., 2023). When using metabarcoding data as a quantitative metric of species relative abundance, some effort must be made to correct for this amplification bias using either internal DNA standards (Sato et al., 2021; Tsuji et al., 2022) or mock communities comprised of known quantities of DNA for the species community found in field samples (i.e., Shelton et al., 2023). Bias-corrected metabarcoding data can then be analyzed using standard ecological approaches and compared to traditional sampling methods.

The general observation in comparative studies that eDNA detects some - but not all - of the same fishes as net-based gear types, leads to the question of what underpins differences in gear selectivity. Put another way, which species traits are being ‘selected’ by eDNA more so than by nets? Patterns of trait-based selectivity have been found across aquatic ecosystems (Aglieri et al., 2021; Mbaru et al., 2020). Less common are studies that examine gear selectivity by species functional traits and include eDNA as one of the gear types (but see Czeglédi et al., 2021; Pont et al., 2021), although some studies have made general observations about functional traits, even when traits were not explicitly tested (i.e., eDNA detected larger pelagic species that beach seines did not; (Saunders et al., 2024). By focusing on functional traits, observations about gear selectivity become more generalizable, as common traits are identifiable across similar types of aquatic ecosystems around the globe. Unlike species identities, traits are linked to ecological functions, and could be applied as indicators of ecosystem function (Henseler and Oesterwind, 2023) or as a way to monitor ecosystem changes (Pereira et al., 2013).

Building on the results from previous studies of gear selectivity and eDNA (He et al., 2022; Saunders et al., 2024), our goal is to address the question of *why* some species are more efficiently detected by different gear types. Further, we examine the species functional traits that underpin differences in gear efficiency. We conducted paired sampling using two different gear types; beach seines and eDNA metabarcoding in three types of nearshore habitats (eelgrass, mixed eelgrass, and understory kelp) in southeast Alaska, USA. We selected nearshore habitats because they are productive ecosystems that maintain complex food webs, confer ecosystem services, and serve as fish nurseries for various species including for commercially important fisheries (Beck et al., 2006; Heck Jr. et al., 2003; Lefcheck et al., 2019). In this study, we 1) compared fish communities across habitats sampled with each gear type, 2) identified which species are associated with which gear type, and then 3) quantified the extent to which each of three traits influences detection probabilities by gear type. The three traits that we selected for analysis are characteristics that might affect eDNA detection: species’ position-in-the-water-column, schooling behavior, and presence of scales/scale-type. Unlike capture-based methods, eDNA is advected from its source and disperses, such that water can contain DNA from species that are currently, or were recently, present, with detection typically decreasing with increasing distance (e.g., Andruszkiewicz et al., 2019; Baetscher et al., 2024; Brasseale et al., 2025). Intuitively, larger amounts of eDNA are produced by more fish (e.g., Ledger et al., 2024), which can also increase detection. Thus, we hypothesized that eDNA detections would be greater for schooling fish and for species with deciduous scales that shed easily, and that eDNA would be less effective at detecting demersal species because water samples were collected at the surface (Fig. 1). Conversely, when sampling with beach seines, we hypothesized that detections of mobile schooling fishes (that escape the seine) would be lower than solitary fishes and that position-in-the-water-column and scale type would not impact detections (Fig. 1).

**Figure 1.**
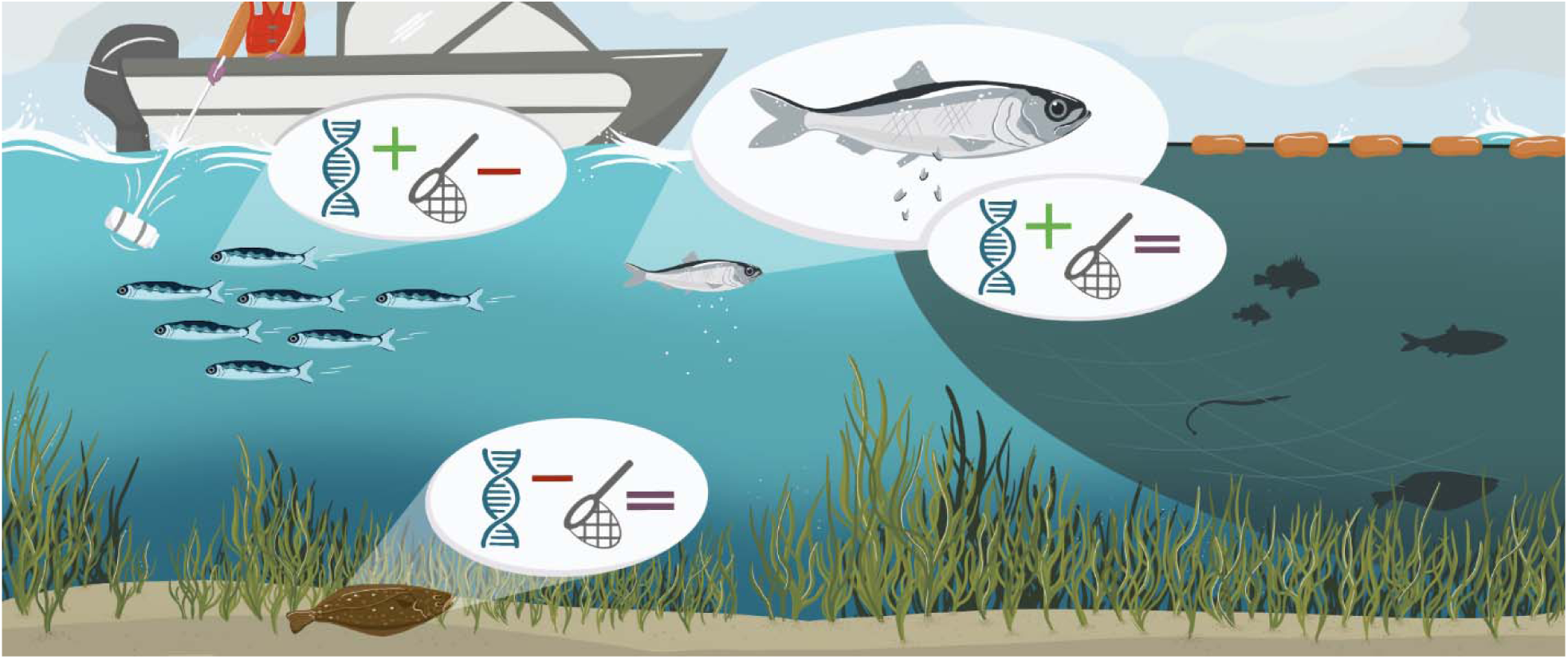
Graphical representation of hypothesized fish species detection based on species traits and sampling method. Hypotheses include: eDNA will more readily detect schooling fish compared to non-schooling fish, whereas beach seines are less likely to detect schooling fish compared to non-schooling fish. Position-in-the-water-column will not influence beach seine detections; however, it will influence eDNA detections with demersal species having lower detection compared to pelagic species. Species that possess deciduous, easily shed scales, will have greater detection with eDNA compared to species with plates or no scales. Scale type will not influence detection with beach seines.

In order to provide a quantitative estimate of the effect of functional traits on gear selectivity, we applied Bayesian occupancy modeling. Occupancy modeling can be used to determine the detection probability of a species because the models account for both species occurrence (i.e., presence, and which variables [e.g., habitat] influence presence) and detection, which includes an evaluation of whether observation methods are suited to detect the species if it is present at the site. This approach allowed us to model the true occupancy of a species based on observations (Doser et al., 2022b). While in many ecological studies, the true species occupancy is the desired outcome, in this study, we apply occupancy modeling to generate a quantitative estimate for the effect of species traits on detection probabilities for both eDNA and beach seines. To our knowledge, the only prior application of occupancy modeling in conjunction with eDNA and gear selectivity is in the context of invasive species (Pukk et al., 2021).

Our study quantifies how functional species traits influence detection by eDNA and traditional sampling methods, which is a critical precursor to accurately characterizing species distributions across a spectrum of species traits. This approach provides a framework for quantifying differences in gear selectivity, particularly for eDNA as it quickly becomes a popular sampling method for understanding species distributions and fish communities.

## 2 Materials and Methods

### 2.1 Study area

We sampled sites in southern Southeast Alaska, around Prince of Wales Island, Revillagigedo Island, and surrounding smaller islands (Fig. 2). This region hosts a diversity of nearshore habitats from low-energy soft-sediment estuaries and protected bays to highly exposed rocky habitats. Low-energy areas and protected bays can have expansive and narrow, fringing eelgrass meadows (*Zostera marina*) in low intertidal and subtidal zones. In cobble and rocky sediment, vegetation shifts to understory kelps (*Laminaria, Saccharina,* and *Hedophyllum* spp. among others), surfgrass (*Phyllospadix serrulatus*), and canopy kelps (*Nereocystis luetkeana* and *Macrocystis pyrifera*). We sampled fish assemblages across a variety of these habitats, including on a gradient from expansive eelgrass meadows (n = 17) to eelgrass meadows with adjacent habitats (understory kelp or canopy kelp) which we call “mixed eelgrass” (n = 11), to understory kelp habitats (n = 7). Candidate sites were identified using a historical and publicly-available aerial imagery and shoreline characterization dataset surveyed between 2004-2010 called ShoreZone (Harper and Morris, 2014) or from previously sampled locations in the Nearshore Fish Atlas of Alaska (Johnson et al., 2012).

**Figure 2.**
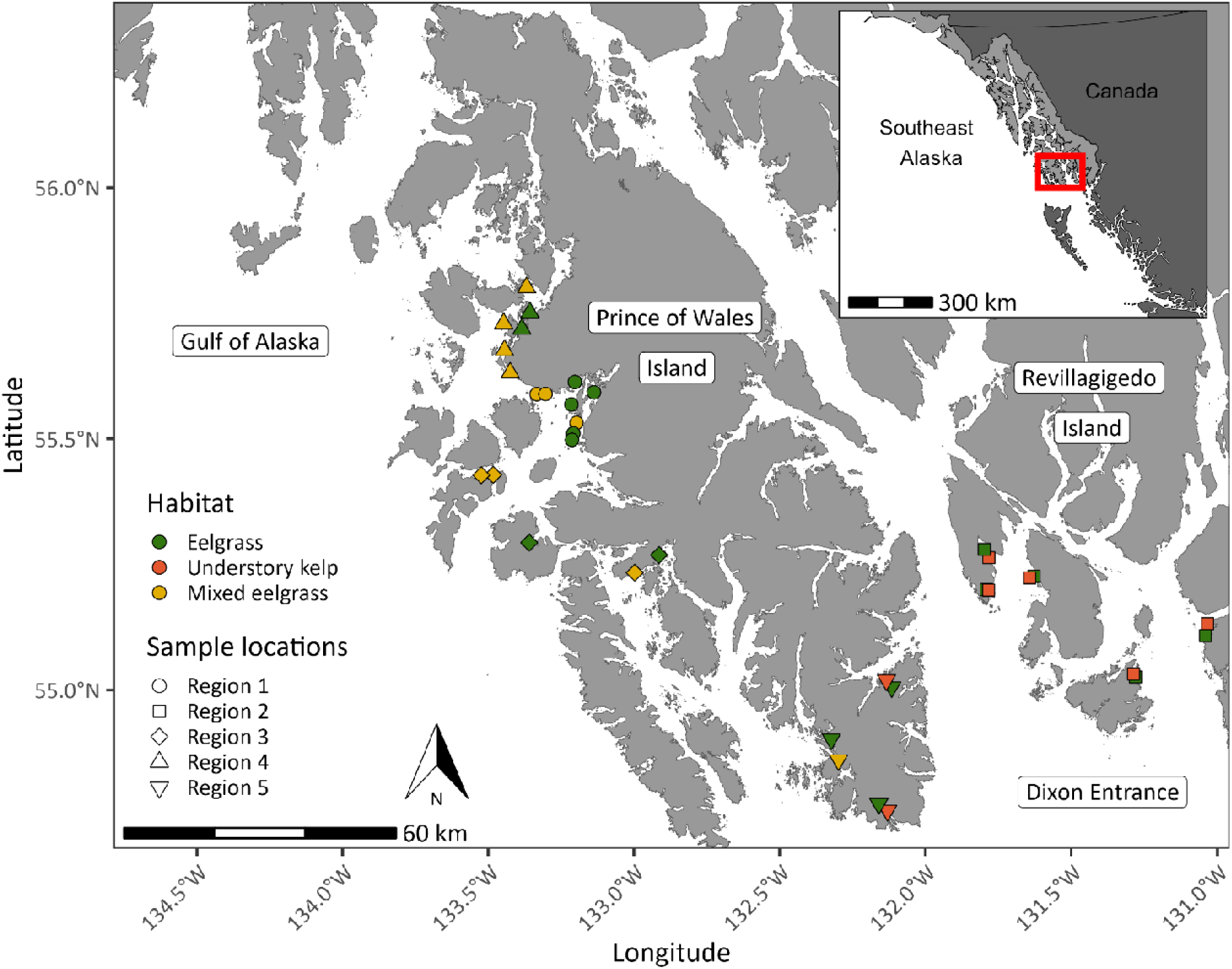
Map of paired eDNA and beach seine sampling locations throughout southeast Alaska, USA. Habitats sampled include eelgrass (green), understory kelp beds (orange), and “mixed eelgrass” meadows where eelgrass is adjacent to understory or canopy kelp (yellow). Sampling locations are grouped regionally (indicated by shape) for occupancy modelling.

### 2.2 Field sampling

#### 2.2.1 Beach seine

To characterize nearshore fish communities in these habitats, we sampled fish using beach seines (as described in Domke et al., 2025; Johnson et al., 2012). Sites were sampled with beach seines in June - July in 2021 and 2022 during negative low-tide cycles (-0.003 to -0.8 m below mean lower low water, MLLW) with a 37 m variable mesh net tapered in width from 10 m in the center to 5 m in the sides with variable mesh, 32 mm square mesh along sides, decreasing to 6 mm square mesh towards the center with a 3.2 mm square mesh in the middle. We set the net by boat in a round haul, brought the net to shore, and identified and counted all fish to the lowest possible taxonomic level. We conducted fish sampling with permits from the Alaska Department of Fish and Game (#CF-21-062 and CF-22-070) and with approval from the Institutional Animal Use and Care Committee at University of Alaska Fairbanks (project #892147).

#### 2.2.2 Environmental DNA

At the same sites where we collected samples with beach seines, we also collected water samples and processed them for eDNA. At each site, we collected 3 replicate 1 L samples approximately 10 m apart above the target vegetation (i.e., eelgrass or understory kelp) within 0.3 m of the surface in 1.5 - 2 m of water. Samples were collected in bleach-sterilized Nalgene bottles, and collection occurred before beach seining and during the dropping tide. After eDNA sampling, a field blank was collected by filling a clean 1 L bottle with distilled water. Each sample was filtered within 3 hours through 0.45 µm cellulose nitrate filters and preserved in 5 mL of Longmire’s buffer at room temperature until DNA extraction (Renshaw et al., 2015).

### 2.3 Laboratory Methods

#### 2.3.1 Mock Community Assembly

We used mock communities (i.e., known mixtures of DNA) to determine the species-specific amplification efficiencies for 22 taxa in this study. We constructed 13 mocks with 7-15 taxa each. With the exception of two species that were included in only one mock, all other taxa were represented in 3-11 mocks (Table S2).

#### 2.3.2 eDNA Extraction, Metabarcoding Amplification, and Sequencing

Detailed eDNA extraction, amplification and sequencing methods are provided in the Supplementary Information. Briefly, eDNA was extracted from filters stored in Longmire’s buffer using the Qiagen DNeasy Blood and Tissue kit. A ∼186bp hypervariable region of mitochondrial DNA on the 12S rRNA gene (MiFish; Miya et al., 2015) was amplified using modified primers from Sales et al. (2019). Each extraction and mock community was amplified in triplicate, and technical PCR replicates were uniquely barcoded to capture stochastic variation during amplification. Library preparation and Illumina MiSeq sequencing of mock communities were performed separately from environmental samples.

### 2.4 Bioinformatic Analysis

Demultiplexed sequences were processed using the Dadasnake pipeline, which implements cutadapt and DADA2 to generate amplicon sequence variants (Callahan et al., 2016; Martin, 2011; Weißbecker et al., 2020). Taxonomic assignments were performed using blastn against the NCBI nucleotide database, followed by custom filtering to exclude ambiguous, non-target, or species outside of the geographic range. ASVs that could not be unambiguously resolved to a single species were consolidated to the highest possible taxonomic level. Additional taxonomic assignment and quality control steps, including correction for tag-jumping and removal of low-read replicates, are described in Supplementary Information.

### 2.5 Data Analyses

#### 2.5.1 Assessing amplification bias in metabarcoding data

We used mock communities to estimate species-specific amplification efficiencies using the quantitative metabarcoding model described in Shelton et al. (2023), which leverages the discrepancy between known starting proportions of DNA and observed proportions of reads after the exponential process of PCR and sequencing. First, we estimated independent amplification efficiencies for each of the 13 mocks in isolation. This resulted in efficiency estimates for each species in each mock community (i.e., if *Clupea pallasii* was in five mock communities, then we generated five efficiency estimates for *C. pallasii*). We applied a centered log-ratio (CLR) transformation to the efficiency estimates and visualized the means and interquartile ranges (IQRs) to identify any mocks with species estimates consistently deviating from the species estimates in the other mocks. Mocks were excluded if two or more taxa had non-overlapping 95% IQRs compared to other mocks (Fig. S2). We then used all of the remaining mocks to jointly estimate a single set of amplification efficiency means and IQRs for each taxa. We calculated and visualized the means and IQRs of CLR transformed efficiency estimates (Fig. S3).

Although multiple factors contribute to amplification bias, primer-template mismatches have the most pronounced effect (Shaffer et al., 2025). To assess these mismatches, we downloaded representative 12S reference sequences for all species detected using beach seine or eDNA sampling. Of these, 54 species had sequences of the MiFish amplicon long enough to include the primer-binding regions. Sequences were trimmed to the primer-binding sites and aligned using Geneious Prime to quantify the number of mismatches for each species.

#### 2.5.2 Influence of gear type and habitat

To quantify the influence of gear type and habitat on nearshore fish community composition, we evaluated differences in species composition using multivariate analysis. Because eDNA metabarcoding data are compositional, we transformed beach seine abundance data to proportional abundance. We applied Wisconsin double standardization on the combined eDNA and beach seine data to minimize differences in absolute abundance across sites and species. Bray-Curtis dissimilarity metrics were then calculated using the vegan package in R (Bray and Curtis, 1957; Jongman et al., 1995; Oksanen et al., 2022). We tested the assumption of homogeneity of dispersion using ‘betadisper’ function in the vegan package in R for subsequent permutation-based tests of differences with gear type and habitat (Anderson, 2006; McArdle and Anderson, 2001; Oksanen et al., 2022). We evaluated differences in species composition between gear type, habitat, and the interaction of gear type and habitat using the vegan ‘adonis2’ function in R, which performs permutation-based analysis of variance (PERMANOVA) and included blocks by site to account for the random effect of multiple eDNA samples per site (Oksanen et al., 2022). For each significant difference (alpha = 0.05) in species composition, we used sequential post hoc pairwise tests to determine which combinations of gear type and habitat were different from each other, focusing only on combinations that represented biological reality. Combinations included a single habitat with different gear types (1: eelgrass/eDNA vs. seine, 2: mixed eelgrass/eDNA vs. seine, 3: kelp/eDNA vs. seine) and the same gear type in different habitats for eDNA (4: eDNA/eelgrass vs. kelp, 5: eDNA/eelgrass vs. mixed eelgrass, 6: eDNA/kelp vs. mixed eelgrass) and beach seines (7: seine/eelgrass vs. kelp, 8: seine/eelgrass vs. mixed eelgrass, 9: seine/kelp vs. mixed eelgrass).

When fish community composition differed significantly by gear, habitat, or their interaction, we used indicator species analysis (ISA) to identify which species were associated with eDNA or beach seine (gear), habitat, or combinations of both. ISA identifies how strongly a species associates with a particular group (or combination of groups) by calculating a species relative abundance (specificity) and relative frequency (fidelity) to a particular group (Dufrêne and Legendre, 1997). The ISA indicator value ranges from 0-1, where 0 indicates a species is absent, and one indicates species that occur only within a specific habitat, gear, or habitat/gear group. Permutation based tests were used to determine significance of species and group associations implemented with the ‘multipatt’ function in the indicspecies package (De Cáceres et al., 2010) with a Bonferroni correction for multiple comparisons. For each species that was significantly associated with a group or combination of groups, we visualized the indicator value magnitude across all logical combinations of gear type and habitat using a heatmap.

In order to investigate the role of species functional traits on gear selectivity, we identified traits for all species identified by either gear type within each trait-category (scale-type, position-in-the-water-column, and schooling behavior; Table S3) using common reference material (ADFG, 2008, n.d.; Eschmeyer and Herald, 1983; Mecklenburg et al., 2002). Each trait-category had several levels. For scale-type, we determined if a fish had no scales, plates, some scales, scales, or deciduous scales. We had three levels for position-in-the-water-column: demersal, pelagic or both demersal and pelagic and three levels for schooling behavior: solitary, facultative schooling, and obligatory schooling. For our multivariate analysis, we created trait groups with all species that shared the same combination of trait-levels within each trait-category. For example, several poachers (*Agonopsis vulsa, Pallasina barbata,* and *Podothecus accipenserinus*) and several sculpins (*Blepsias bilobus, Blepsias cirrhosis, Nautichthys oculofasciatus,* and *Rhamphocottus richardsonii*) represented the trait group of solitary, demersal, and plated fishes. We visualized any patterns in species compositions by gear type and habitat with a principal coordinate analysis (PCoA) and plotted the combined trait-category groups (described above) that significantly correlated with each axis.

To evaluate the impact of species functional traits on detection probability by gear type, we used Bayesian integrated occupancy models in the ‘spOccupancy’ R package (Doser et al., 2022b, 2022a; Miller et al., 2019). First, we transformed our eDNA and beach seine relative-abundance data to binary detection/non-detection data for all 70 species detected by one or both gear types. Occupancy models leverage repeated observations of detection/non-detection data to jointly estimate detectability and probability of occurrence (MacKenzie et al., 2002). In our integrated approach, the two gear types produced datasets with separate observation process models linked together by a shared site-level occupancy process. Whereas occupancy models are typically used to facilitate inference on occupancy by accounting for imperfect observations, our interest was to use the models to facilitate inference on detection by accounting for heterogeneous patterns of species’ presence. In the multispecies context, we assumed species-specific coefficients were random effects that arise from common distributions with community-level mean and variance terms. We hypothesized that species’ probabilities of occurrence or use at a site was influenced by habitat and therefore included a site-specific covariate for habitat (based on results from pairwise tests; Table S4) as well as a site-specific covariate for region (Fig. 2), as sites that were closer together may have had more similar species composition. We assumed a constant probability of detection for both gear types and thus fit intercept-only detection submodels for each gear type. Model convergence was assessed using the Gelman-Rubin diagnostic (Rhat; Brooks and Gelman, 1998) and effective sample size of the posterior samples to ensure adequate mixing. We evaluated model fit using posterior predictive checks following Doser et al. (2022b), by calculating Bayesian p-values based on Chi-square discrepancy statistics calculated for observed and predicted data binned across replicates (code provided in Supplementary Information).

To determine if species’ functional traits influenced detection by gear type, we fit post hoc secondary linear models (sometimes referred to as a 2-stage analysis), in which the species-specific logit-scale detection parameters from each iteration of the posterior of the occupancy model were treated as response variables (Doser et al., 2022a). This was done separately for each gear type, allowing us to investigate the influence of species functional traits on the probability of gear-specific detection. We interpreted effect sizes on the odds-ratio (OR) scale. Odds ratios, calculated as exp(logit-scale coefficient) for each trait, lie on a strictly positive scale, where OR>1 indicated a greater odds of detection relative to the reference trait combination (positive association), OR<1 being a negative association, and a OR=1 being no association between the trait and detection probability. We interpreted the proportion of each posterior distribution that fell above zero on the logit scale (or above or below 1 on the odds-ratio scale), as the probability of a positive association between the corresponding trait and detection probability relative to the reference trait (or the probability of a negative association for the proportion below zero). Thus, if 95% of a logit-scale posterior was >0, that indicated a probability 0.95 of a positive association (and 0.05 for a negative association) between the corresponding trait and detection probability.

## 3 Results

After performing quality control on sequence reads from environmental and mock community samples, the final dataset included 4.83 million reads from 576 ASVs representing 66 taxa in field samples, and 1.46 million reads from 50 ASVs representing 24 taxa in mock communities (see Supplementary Information for full details).

### 3.1 Lack of Consistent Amplification Bias

One of the 13 mock communities had three taxa with non-overlapping IQRs compared to the other mocks and was therefore excluded from subsequent analyses (Fig. S2). Excluding this mock removed 2 of the 22 species present across all 13 mocks, leaving 20 species for which amplification efficiency was calculated. Among those 20 species, there were no differences in amplification efficiency based on IQRs of centered-log ratio (CLR) amplification estimates (Fig. S3).

As a potential source of amplification bias, we examined primer-template mismatches. Of the 54 species with reference sequences spanning the primer sites, three taxa had a single mismatch and two taxa had two mismatches. No species in the mock communities contained primer-template mismatches (Fig. S4). Given that estimated amplification efficiencies across species in the mock communities were relatively similar and few species contained primer mismatches, we did not apply the quantitative metabarcoding model to field samples. Because our dataset did not require using the quantitative metabarcoding model, we could expand the number of species analyzed beyond those in the mock communities to include all detected taxa and calculated simple means of metabarcoding read proportions from the technical replicates.

### 3.2 Gear Type and Habitat Influence Observed Fish Community Composition

Overall, eDNA and beach seines identified 70 species (eDNA = 66, seine = 49) with four species only detected with seines and 21 species only detected with eDNA. Across the three habitat types eDNA identified more species than beach seines in each habitat (Fig. S5; eDNA: 53 species [eelgrass], 50 [mixed eelgrass], 48 [understory kelp]; seine: 38 [eelgrass], 42 [mixed eelgrass], 34 [understory kelp]).

Fish community composition differed significantly with the interaction of gear and habitat (PERMANOVA, Fig. 3, Table 1). We detected no within-group dispersion for gear type (F-statistic_1, 135_ = 1.91, p-value = 0.169) or habitat (F-statistic_2, 134_ = 1.89, p-value = 0.154). We found strong evidence for differences in species composition in each of the nine logical pairwise combinations of gear type and habitat (Table S4, p-values < 0.05). We visualized species composition with principal coordinates analysis (PCoA) with the interaction of gear and habitat type representing 40.4% of the total variability in species composition with the first two axes (Fig. 3).

**Figure 3.**
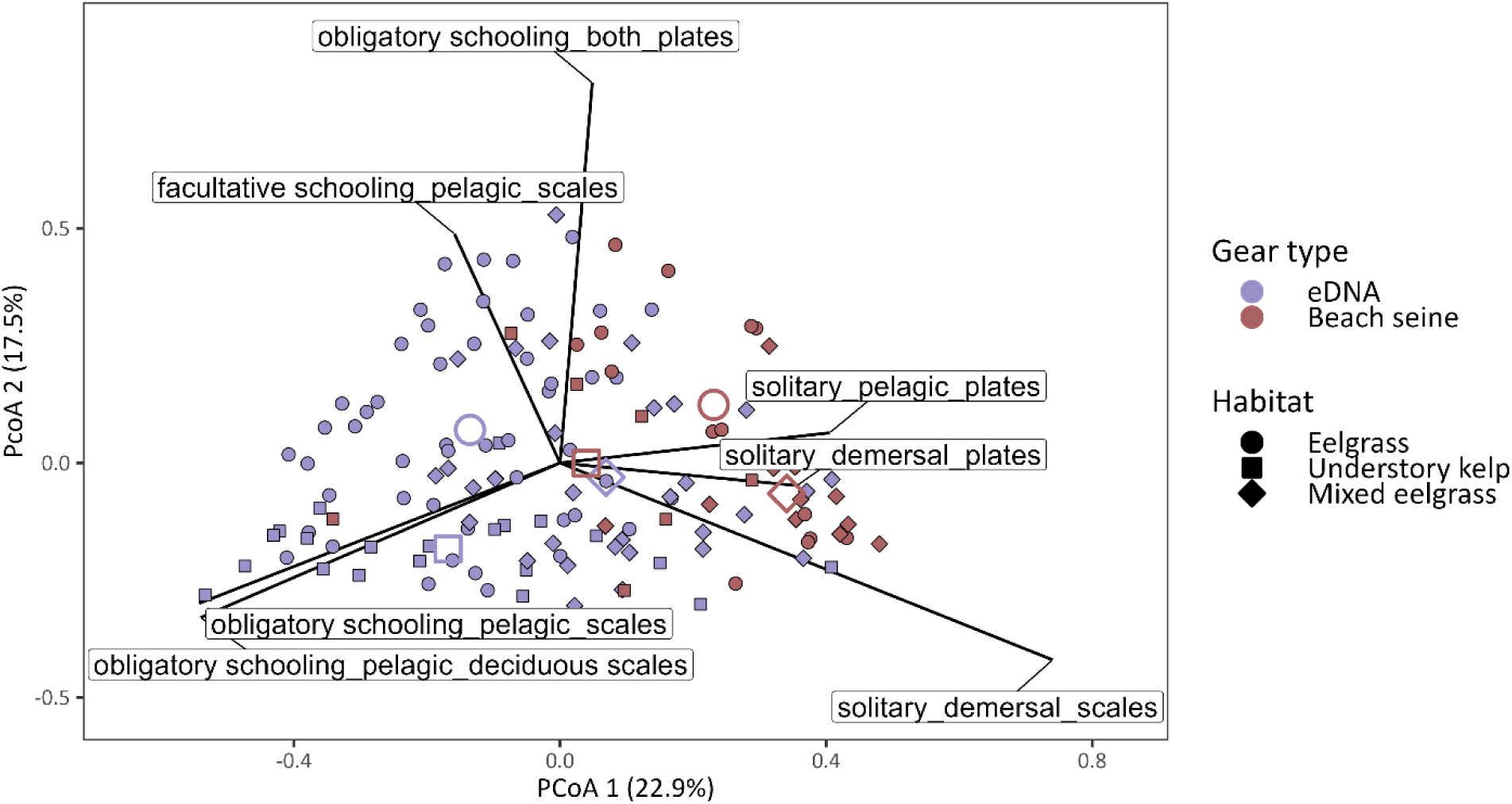
Association of gear types with traits. Principal coordinate analysis (PCoA) of species composition sampled by eDNA (purple) and beach seine (red) in eelgrass (circle), mixed eelgrass (square), and understory kelp (diamond). Each point represents the composition of fish species for a single replicate at a given site, with unfilled larger points representing the average position of points for one gear type-habitat combination. Vectors indicate trait combinations (e.g., schooling behavior, position-in-water-column, and scale type) that are significantly correlated with PCoA axes (p-value < 0.01), with the length and direction of the lines indicating the strength and direction of the relationship. For example, obligatory schooling species that are pelagic with scales or deciduous scales are more associated with the lower left quadrant of the PCoA where fish communities sampled with eDNA are most prevalent. Biological replicates for eDNA samples taken at the same site are shown, whereas only one sample was collected at each site by the beach seine.

**Table 1.** Permutation-based ANOVA (PERMANOVA) of the interaction between habitat type (eelgrass, mixed eelgrass, and understory kelp) and gear type (eDNA and beach seine) for fish community composition. Permutations were performed with blocks to account for within site variability across multiple eDNA water samples collected at each site.

| | DF | Marginal $R^2$ | Sum of<br>squares | Pseudo-F<br>statistic | P-value |
| --- | --- | --- | --- | --- | --- |
| Habitat × Gear type | 2 | 0.026 | 1.207 | 1.996 | 0.002 |
| Residual | 131 | 0.845 | 39.586 |  |  |
| Total | 136 | 1.0 | 46.870 |  |  |

### 3.3 Indicator Taxa Associated with Gear and/or Habitat

Indicator species analysis (ISA) identified significant associations of 1-5 species with eight gear-habitat combinations, with more indicator species with beach seines compared to eDNA (eDNA mean = 0.7 species/habitat combination, seine mean = 1.7 species/habitat combination; Fig. 4). ISA identified three species with significant indicator values across all habitats sampled with eDNA (*Anoplarchus* sp., *C. pallasii*, and *Oligocottus maculosus*; Fig. 4). Other indicator species for habitats sampled with eDNA included *Xiphister* in understory kelp.

**Figure 4.**
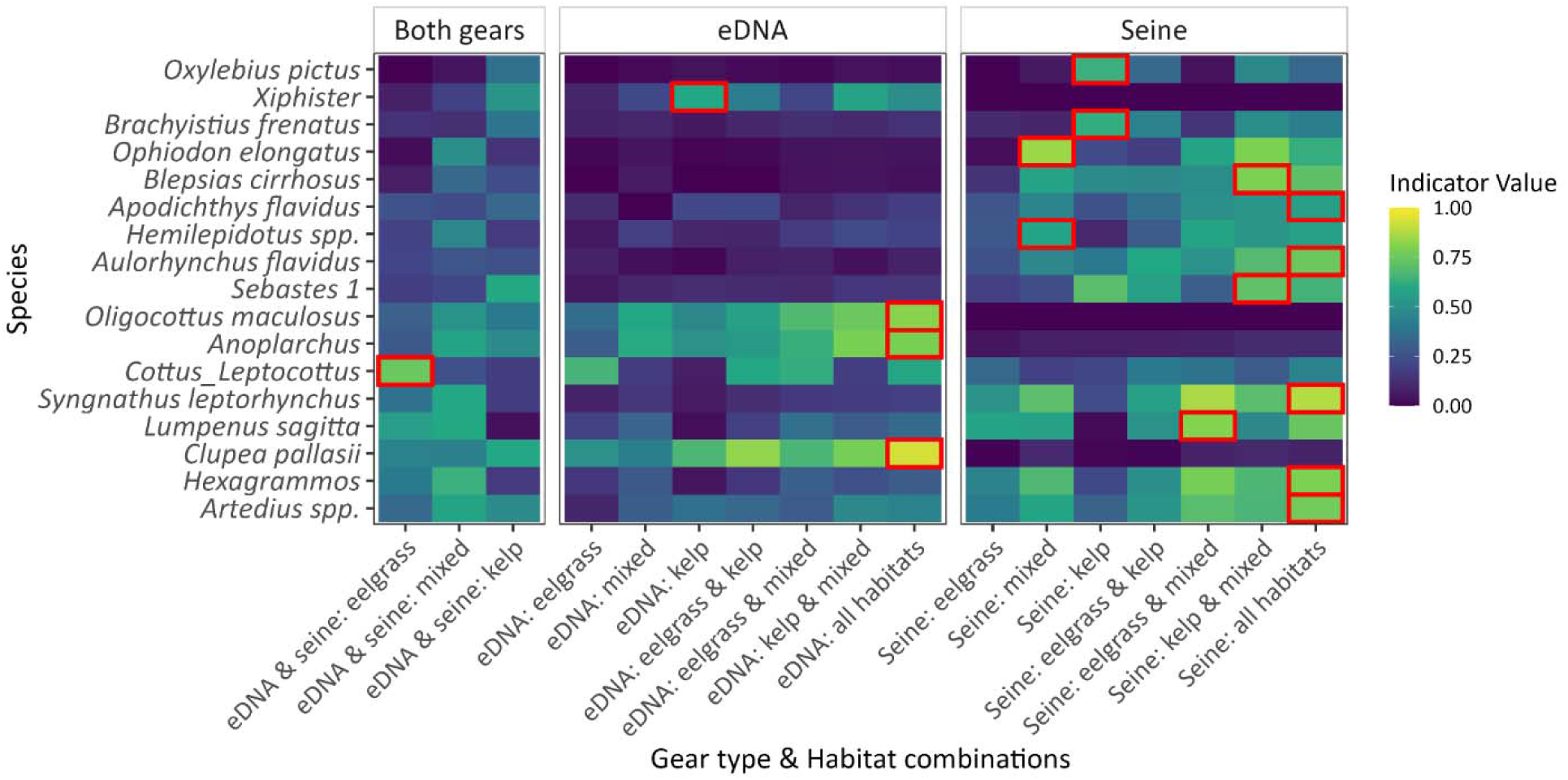
Indicator species are associated with gear type and habitat combinations. Indicator values for species with a significant association for a given combination of gear types (eDNA, seine, or both) and habitats (eelgrass, kelp, mixed) are boxed in red (Bonferroni corrected p-value < 0.05). Indicator value ranges from 0 (dark blue), where a species is absent, to 1 (yellow), where a species occurs exclusively with a gear/habitat combination. For example, *Cottus_Leptocottus* (a group of sculpins that are taxonomically indistinguishable with the 12S MiFish metabarcoding marker) is an indicator species for eelgrass sampled with eDNA or seine because both gear types captured relatively high abundance and fidelity of this species in eelgrass, and not in other habitats.

Indicator species varied by gear type. Five species were indicator species across all beach-seined habitats (*Artedius* spp., *Hexagrammos* spp., *Syngnathus leptorhynchus*, *Aulorhynchus flavidus*, and *Apodichthys flavidus*). Indicator species for understory kelp and mixed habitats included *Sebastes* 1 (see Table S2) and *B. cirrhosus*. *Lumpenus sagitta* was the sole indicator species for eelgrass and mixed eelgrass. For beach seining in understory kelp sites, we identified two indicator species: *Brachyistius frenatus* and *Oxylebius pictus,* and in mixed eelgrass we identified two indicator species, *Ophiodon elongatus* and *Hemilepidotus* spp. The only indicator species significant in both gear types was the group with the genera *Cottus* and *Leptocottus* in eelgrass habitats (Fig. 4).

### 3.4 Species traits influence fish community sampled

#### 3.4.1 Trait-Based Assemblages across Gear Types and Habitats

Seven trait group vectors were correlated with the PCoA axes (p-value < 0.01) with the length and direction of the lines indicating the strength and direction of the relationship (Fig. 4). These significant trait groups included 1) obligatory schooling species that inhabit both demersal/pelagic positions-in-the-water-column and have plates rather than scales, 2) facultative schooling species that are pelagic and have scales, 3) solitary pelagic species with plates, 4) solitary demersal species with plates, 5) solitary demersal species with scales, 6) obligatory schooling pelagic species with scales, and 7) obligatory schooling pelagic species with deciduous scales.

#### 3.4.2 (Some) Functional Traits Drive Variation in Detection Probability by Gear Type

We observed different detection probability by gear type and species’ functional traits. Species with a combination of scales, solitary, and demersal traits (represented by the intercept) had similar detection probability with both eDNA and beach seines (mean and 95% credible interval for detection probability = 0.216 [0.12-0.34; eDNA] and 0.211 [0.09-0.40; seines]; Fig. 5). We found evidence of associations between trait-levels and gear-specific detectability for three trait-gear combinations (using the criteria that ≥95% of the covariate’s odds ratio posterior distribution fell above or below 1). With eDNA – but not beach seines – detection probability for a species with deciduous scales was greater than that of otherwise similar species that shared all other traits: a species with deciduous scales had on average 27 times greater odds of detection compared to a similar species with scales (p(OR>1) = 0.97). Conversely, a fish with plates rather than scales had lower odds of detection with eDNA (0.31 times lower compared to a similar fish with scales; p(OR>1) = 0.04), but showed some indication of greater odds of beach seine detection (again, relative to scaled fish) with 2.4 times greater odds (p(OR>1) = 0.81). Fish with no scales had reduced detection probability with beach seines compared to fish with scales (p(OR>1) = 0.05), while this trait was not associated with any difference in detection with eDNA (p(OR>1) = 0.53). Fishes that were both demersal and pelagic compared to just demersal had increased detection probability with both eDNA and beach seines, with 3.9 and 19.8 times greater odds of detection, respectively (p(OR>1) = 0.77 [eDNA] and p(OR>1)=0.86 [seines]). For several traits there was no clear evidence of difference in detection probability by trait-levels in either gear type, including presence of some scales, obligatory and facultative schooling traits, and pelagic species.

**Figure 5.**
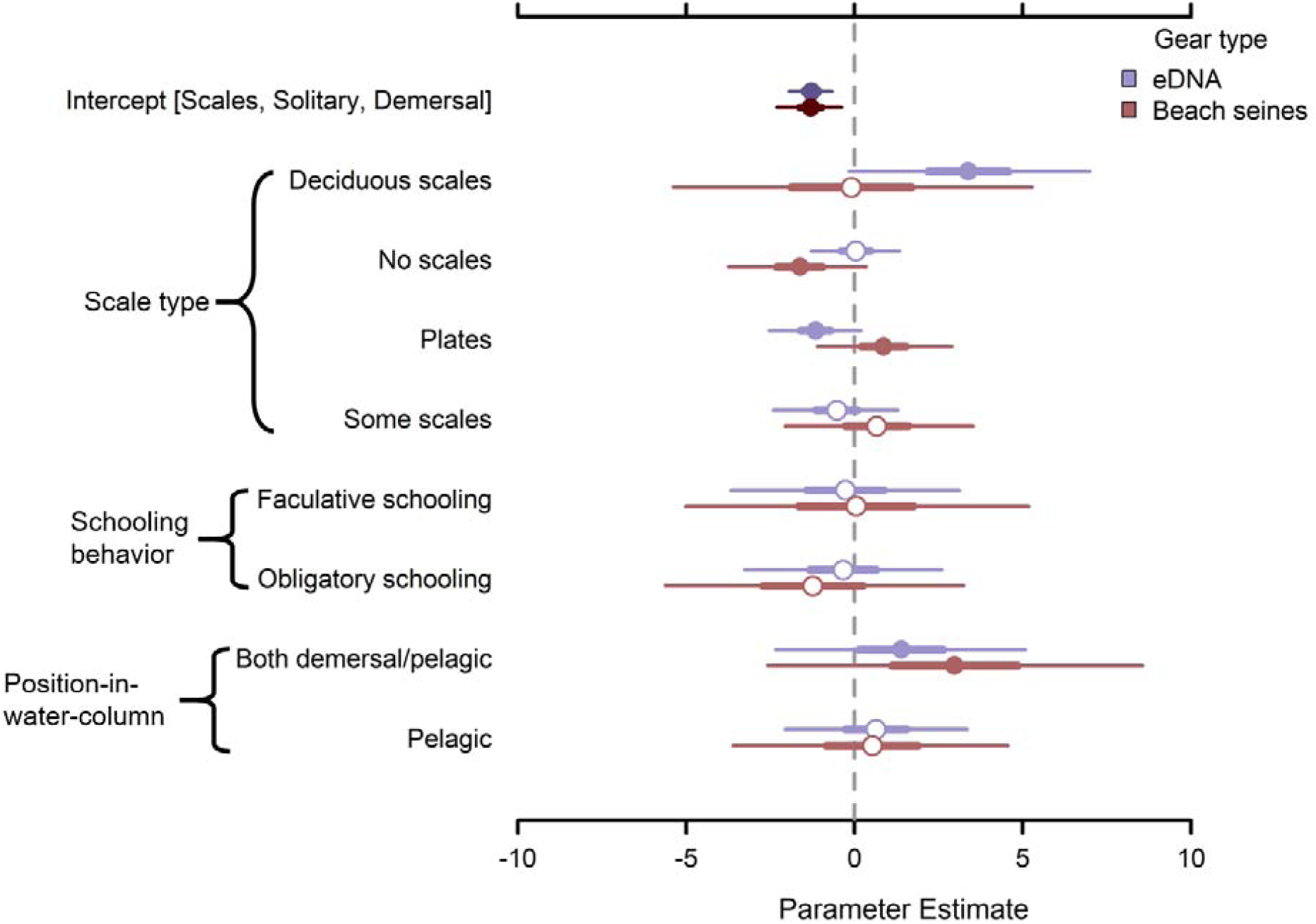
Median trait effects on detection by gear type from post-hoc secondary models. Logit-scale estimates indicate influence of functional trait on detection probability for each gear type: eDNA (purple) or beach seines (red) relative to the reference trait combination (i.e., the intercept, represented by a fish that has scales, is solitary, and demersal). Thick and thin bars correspond to 50% and 95% credible intervals (CI); filled circles indicate that the 50% CI does not overlap 0. Each gear type was modeled independently. For example, a fish with plates that is solitary and demersal will have decreased detection with eDNA and increased detection with beach seines relative to a fish that has scales and is solitary and demersal.

## Discussion

Studies often make ecological inferences based on sampling community composition, but each sampling method has selectivity biases that lead to some species being more readily detected. In this study, we identified the functional traits underlying selectivity of eDNA and beach seines in fishes across three nearshore habitats in southeast Alaska. We found significant differences in the fish community across eelgrass meadows, understory kelp, and mixed eelgrass habitats when sampled with eDNA and beach seines. Fewer than 20 species out of the 70 detected were identified as indicator species for combinations of habitat and gear type. Functional traits hypothesized to influence differences in gear selectivity between eDNA and seines included a species’ position-in-the-water-column, schooling behavior, and scale-type/presence of scales. Of these traits, scale-type had the greatest impact on probability of detection by either gear type. While gear selectivity for specific functional traits has been qualitatively inferred in previous studies, our use of occupancy models to quantify differences in detection probabilities by gear type is a novel extension of previous analyses.

Of the 70 species detected by either eDNA or beach seine, 17 species (or species groups) were identified in the indicator analysis as being associated with one or more gear-types and habitats (Fig. 4). Only one species group, *Cottus Leptocottus* (a group of two genera of sculpins that are taxonomically indistinguishable with the MiFish metabarcoding marker), was significantly associated with both gear types, and only in eelgrass meadows. The strongest indicator value for any species across all gear-type/habitat combinations was *Clupea pallasii*, Pacific herring, which was significantly associated with eDNA sampled in all habitats. This result was consistent with our hypotheses about the ability for eDNA to detect schooling species with deciduous scales more readily than seines (Fig. 1). Otherwise, patterns of association were difficult to disentangle between functional traits, species, and habitats.

Because indicator species analysis takes into account both relative abundance and fidelity, species that are not abundant, but have high fidelity, may still be significant. *O. pictus* (painted greenling), for example, were only significant in kelp habitat sampled by seines. Since *O. pictus* has scales (but not deciduous scales), it is possible that low relative abundance (Fig. S6), coupled with solitary and demersal traits, kept the species from receiving a higher proportion of eDNA sequencing reads. Seines may have yielded more indicator species than eDNA because their method of capture provides a precise link between species and habitats. In contrast, the dispersal of eDNA creates a “smoothing effect” on species detections which may make it a less precise indicator of fine-scale habitat use. Further, multiple eDNA water samples were collected at each site, potentially increasing the likelihood of detecting nearby species.

The PCoA’s primary axis of variation separated the communities detected by eDNA (left) from those detected by seines (right). This axis also separated the functional traits of schooling behavior and scale-type. While schooling behavior was an inconclusive functional trait based on the indicator species analysis, the primary axis of the PCoA separated trait groups that included schooling species from solitary species (Fig. 3), which suggested that eDNA preferentially detected schooling species. Additionally, trait groups that included scales or deciduous scales (with the exception of the solitary-demersal-scale group) separated along this axis, suggesting a relationship of improved detection with eDNA compared to scales. No clear pattern was found linking species’ position-in-the-water column to the PCoA axes. Ultimately, the relationship between gear detection and functional trait is potentially confounded by multiple interactions involving habitat, species distribution, and trait expression (Fig. 3).

Trait-based quantitative occupancy modeling allowed us to disentangle the potential for gear selectivity on multiple interacting traits. A prime example of this phenomenon is that increased detection probability of *C. pallasii* when sampled with eDNA could have been caused by deciduous scales, schooling behavior, or an unattributed trait. Based on results from the Bayesian trait analysis, species with deciduous scales were more likely to be detected by eDNA than species with non-deciduous scales (Fig. 5). In comparison, schooling behavior did not appear to impact detection probabilities for either gear type. The impact of scale type on detections can be observed in comparing detections of *C. pallasii* (deciduous scales) and *S. leptorhynchus* (plated). We observed *C. pallasii* in nearly all our sites with eDNA and rarely with beach seines and the opposite pattern with *S. leptorhynchus*, which was caught in most sites with beach seines, but rarely detected with eDNA (Fig. S6).

Traits other than the ones we evaluated (impact of schooling behavior, position-in-the-water-column, and scale type), could influence gear selectivity. Previous studies of gear selectivity in nearshore habitats have examined traits including diet, body size, depth, swimming speed, and period of activity, in addition to schooling behavior and position-in-the-water-column (e.g., Henseler and Oesterwind, 2023; Mbaru et al., 2020). While many studies identify differences between functional traits detected by different gear types, most of these studies use a combination of traditional fishing and sampling gears (e.g., Carvalho et al., 2021; Mbaru et al., 2020, but see Czeglédi et al., 2021 for a comparison between eDNA, electrofishing, and gillnetting). Consequently, some traits analyzed in other studies may be less likely to impact detection probabilities for eDNA sampling (e.g., feeding behaviors; Franco et al., 2012; Henseler and Oesterwind, 2023). Investigating certain traits can impact how effective a gear type appears. For example, Aglieri et al. (2021) found that eDNA identified the full spectrum of functional traits considered in their study; however, untested traits, like scale type, could have a confounding effect on detection with eDNA. Likely additional ecological and functional traits are relevant to evaluating nearshore fish communities, in particular with eDNA, which may not be captured in the present study. Future studies evaluating traits influencing eDNA detections may consider including attributes related to DNA shedding, metabolic, and/or movement rates (Ely et al., 2021).

Using metabarcoding data as a reflection of relative species composition requires understanding the impact of amplification bias on the dataset (Shelton et al., 2023). The primary conclusion from our mock community data was that none of the species that we tested exhibited consistent and significant amplification bias. Given the substantial amount of effort required to construct mock communities, finding a method to test for potential amplification issues prior to beginning lab work would be valuable. For example, if amplification bias is primarily driven by primer-template mismatches (i.e., Shaffer et al., 2025), then in silico sequence alignments could provide valuable information as to whether mock communities are required to correct amplification biases in metabarcoding data (e.g., Fig. S4). Even if primer-template mismatches are present, their impact may be trivial if all species have similar numbers or types of mismatches because sequencing read proportions are dependent on the relative amplification efficiencies across all species in a sample (Shaffer et al., 2025). Five species had primer-template mismatches in the fish communities sampled in this study (Fig. S4). Two of those species *Mallotus villosus* (capelin) and *Podothecus accipenserinus* (sturgeon poacher) were absent from eDNA sampled at sites where they were present in beach seines. Given the small number of mismatches (1-2 bases) and their location toward the middle of the primer (Lefever et al., 2013), other sources of sampling variation could just as easily account for observed differences between eDNA and seine detections.

Taxonomic assignment is the cause of much angst in eDNA metabarcoding studies (Blackman et al., 2023; Locatelli et al., 2020). Here, we aggregate uncertain species-level assignments to multi-species groups within a single genus (e.g., *Sebastes* groups 1-3) or, in some cases, a family (e.g., Pleuronectidae). The same taxonomic uncertainty can also occur in beach seine sampling based on potential misidentification, especially of small and hard to distinguish taxa (e.g., *Artedius* spp. sculpins). Despite the potential for misidentification, we doubt the aggregation of species to genus or family impacts the analysis of gear selectivity given the similarity of the functional traits that we chose within taxonomic groups.

Increasingly, multi-gear assessments could mitigate sampling biases of individual gear types rather than benchmarking one gear (typically eDNA) against another (most often traditional sampling; i.e., nets). In fact, multiple studies found that a combination of two sampling methods can adequately sample fish communities (French et al., 2021; Henseler and Oesterwind, 2023) assuming the gear types are complementary (Andres et al., 2023; Stat et al., 2019). In our study, the clearest example of this was the selectivity of eDNA for deciduous scales that allowed for *C. pallasii* to be identified across habitats where it was entirely absent in the beach seines. The framework we describe for analyzing the effect of functional traits on gear detection probabilities provides a model for how similar quantitative assessments could be performed in future studies. Additionally, data from multi-gear assessments could be used to build joint models, which can integrate information from multiple gear types in a spatially explicit framework (e.g., Guri et al., 2024; Keller and Kelly, 2025 to estimate abundance/biomass or Doser et al., 2022a for occupancy presence/absence). Study design and choice of sampling gear should align with which species (and which functional traits) are being targeted. eDNA represents a particularly valuable addition to sampling gears because it is relatively non-invasive, can be used in areas restricted to other types of sampling (i.e., marine reserves), and likely has reduced impacts to species and ecosystems when compared to nets.

## Acknowledgements

Thanks to E. Antaya, E. Beaver, A. E. Kelsey, E. Vernon, I. Sears, and C. J. Johnson who participated in field sampling in 2021 and 2022. O. Shelton provided helpful discussion about the analysis. This research took place on the land of the Tlingit and Haida who continue to steward the land and its resources.

## Author contributions statement

LKD: Conceptualization, Data curation, Formal analysis, Investigation, Methodology, Visualization, Writing – original draft

KJL: Conceptualization, Data curation, Formal analysis, Methodology, Writing – original draft

JMW: Data curation, Investigation, Methodology, Visualization, Writing – review & editing

SMK: Conceptualization, Methodology, Formal analysis, Software, Writing – review & editing

WL: Conceptualization, Methodology, Project administration, Resources, Supervision, Writing – review & editing

GLE: Conceptualization, Funding acquisition, Investigation, Methodology, Project administration, Supervision, Writing – review & editing

DSB: Conceptualization, Methodology, Supervision, Writing – original draft

## Data Availability Statement

Lia K. Domke & Shannon M. Kachel. 2025. NearshoreMiFisheDNA [dataset]. GitHub. https://github.com/lkdomke/NearshoreMiFisheDNA

Kimberly J. Ledger. 2025. nearshore_eDNA [dataset]. GitHub. https://github.com/kjledger-NOAA/nearshore_eDNA

Data are available through Github for analyses and beach seine data (Domke & Kachel, 2025) and processed eDNA data (Ledger, 2025). Raw metabarcoding data will be available through NCBI SRA upon publication.

