## Supplemental Information for "A case for multi-gear assessments: Detection probabilities of nearshore fish with eDNA and seine nets vary by functional traits"

### Supplementary Information

#### Methods

##### Mock Community Assembly

We used mock communities (i.e., known mixtures of DNA) to determine the species-specific amplification efficiencies for 22 taxa in this study. We constructed mock communities using synthetic DNA fragments (gBlock Gene Fragments, Integrated DNA Technologies, Inc., Coralville, IA, USA) that encompass the 12S rRNA gene region used in this study. DNA fragments were diluted to approximately 0.2 ng/uL and quantified using fluorescence of double-stranded DNA (with the Qubit dsDNA High Sensitivity Quantification kit, Invitrogen). We constructed 13 mocks with 7-15 taxa each. With the expectation of two species that appeared in only one mock, all taxa were represented in at 3-11 mocks (Table S2).

##### eDNA Extraction

eDNA extractions were performed in a dedicated pre-PCR laboratory where all instruments were routinely decontaminated with UV and bleach sterilization. DNA was extracted using the Qiagen DNeasy Blood and Tissue kit modified for eDNA filters stored in Longmire’s buffer. Modifications included a starting input of 400 μL of Longmire's buffer and a final elution volume of 100 μL for each sample (following Baetscher et al., 2024). Three and ten extraction blanks were included with samples collected in 2021 and 2022, respectively. Extracts were stored at -20°C until PCR amplification.

##### PCR amplification

We amplified a hypervariable region of the mitochondrial DNA 12S rRNA gene using primers from Miya et al. (2015) and Sales et al. (2019): MiFish-U-F_mod (5’ *CGACAGGTTCAGAGTTCTACAGTCCGACGATC*GCCGGTAAAACTCGTGCCAGC 3’) and MiFish-U-R (5’ *GTGACTGGAGTTCAGACGTGTGCTCTTCCGATCT*CATAGTGGGGTATCTAATCCCAGTTTG 3’), with Illumina Nextera adapters shown in italics. eDNA extractions, negative controls (sterile water), and positive controls (*Acipenser fulvescens* DNA) were amplified in triplicate using GeneAmp PCR System 9700 Thermocyclers (Applied Biosystems). PCR reactions (12  μL) contained 2x Qiagen Multiplex Mastermix, 200 nM of each primer, and 2 μL of DNA extract. Thermocycling conditions were 95°C for 10 min, followed by 35 cycles of 95°C for 30 s, 60°C for 30 s, and 72°C for 30 s, with a final elongation of 68°C for 10 min. PCR products were visualized on a 2% E-Gel agarose gel (Invitrogen) to confirm amplification of the correct product size.

Unique barcodes were added in a second PCR reaction (10 μL) consisting of 2x Qiagen Multiplex Mastermix, 100 nM of each index, and 2 μL of a 1:5 dilution of the first PCR product. Thermocycling conditions were 95°C for 10 min, followed by 10 cycles of 95°C for 30 s, 65°C for 30 s, and 72°C for 30 s, with a final elongation of 72°C for 5 min. Barcoded products were again checked using E-Gels.

Following amplification and barcoding, samples were quantified using the Quant-iT PicoGreen dsDNA Assay Kit (Invitrogen), normalized to 5 ng/μL, and pooled in equal volumes. Size selection targeted the ~350 bp amplicon using double-sided AMPure XP (Beckman Coulter) bead size selection (bead:library ratios = 0.5× followed by 0.7×). Final libraries were quantified using the QuBit 1× dsDNA HS Assay Kit (Invitrogen), and fragment size distributions were determined using an Agilent Technologies 4200 TapeStation. Libraries were denatured and diluted to 10pM, spiked with 20% PhiX to compensate for the low diversity of the library, and sequenced on an Illumina MiSeq with v2 2×150 paired-end chemistry. In total, three PCR replicates of 187 field samples, positive and negative controls were sequenced on a single Illumina MiSeq run. Mock community libraries were sequenced across two MiSeq runs and kept separate from environmental samples.

##### Bioinformatics

Demultiplexed sequences generated from environmental samples and mock community samples were filtered using Dadasnake, a Snakemake implementation of cutadapt and DADA2 to process amplicon sequencing data (Callahan et al., 2016; Martin, 2011; Weißbecker et al., 2020). The pipeline includes primer removal, quality filtering, sequence trimming, merging, and chimera removal and outputs a sample of amplicon sequence variants (ASVs) for each technical PCR replicate. We performed taxonomic assignment of the ASVs using blastn (Altschul et al., 1990) against the NCBI nucleotide database (access date: 18 July 2024) using 96% sequence identity and 98% query coverage. We then used custom filtering of blastn output to remove matches to ambiguous species assignments (i.e., hybrids, “sp.”) and non-Actinopterygii (bony fish). We also removed matches to fish species that do not occur within our study region (Northeast Pacific) based on the FishBase species distribution repository using the rfishbase package in R (Boettiger et al., 2012). For ASVs with taxonomic matches of >99% sequence identity, we retained matches within the top 0.5%. For ASVs without taxonomic matches of >99% sequence identity, we retained matches within 1% of the top match. For ASVs with a single taxonomic match, we retained matches with at least 98% sequence identity.

When ASVs were not unambiguously assigned to a single species, we consolidated ASVs into the highest resolution taxonomic group. In cases where multiple ASVs matched the same genus or family group, we added a numeric identifier to the genus or family to differentiate the distinct species groups (i.e. ASVs matching *Sebastes caurinus* and *Sebastes maliger* are represented by *Sebastes* 1 and ASVs matching *Sebastes auriculatus* and *Sebastes rastrelliger* are represented by *Sebastes* 2; see Table S1 for full breakdown of taxonomic groups).

Additional bioinformatic quality controls included (1) estimating rates of tag-jumping based on the occurrence of ASVs in positive controls (per sequencing run) and subtracting the maximum proportion of an ASV in the positive controls from all PCR replicates, (2) removal of ASVs without taxonomic assignment, (3) removal of ASVs without any reads in the field samples, and (4) removal of PCR replicates with fewer than 1000 total reads (a threshold determined by examining sequence efforts using species accumulation curves with rarecurve function from vegan package in R (Oksanen et al., 2022; Fig. S1).

#### Results

##### eDNA Metabarcoding

We generated approximately 9.5 million sequencing reads from environmental samples and 1.8 million sequencing reads from mock communities. After preliminary filtering using Dadasnake, we retained approximately 5.4 million reads from environmental samples and 1.5 million reads from mock communities representing a total of 859 ASVs. We detected less than 0.3% tag-jumping per ASV in our positive control replicates and subtracted those read proportions from each corresponding ASV in all replicates. After taxonomic assignment and custom filtering to retain only teleost fishes known to occur in the study area (i.e., the Pacific Ocean), removing ASVs found only in controls, and removing replicates with fewer than 1,000 reads, 586 ASVs remained.

After quality control, no reads remained in PCR negative controls (0/12) or extraction negatives (0/18). Reads were detected in 30 of 82 field blank replicates and 321 of 348 field sample replicates. All mock community replicates (39/39) yielded reads. In our final curated dataset, the mocks contained 50 ASVs representing 24 taxa and 1.46 million reads. Two taxa not intentionally included in the mocks were detected at low levels: *Oncorhynchus* and *Pleuronectidae* 3. *Oncorhynchus* was represented by 50 reads across four replicates and two mocks, accounting for less than 0.0003 of the total reads in those communities. *Pleuronectidae* 3 was represented by 832 reads across 13 replicates and six mocks, comprising less than 0.002 of the total reads in those communities. In our final curated dataset, field samples contained 576 ASVs representing 66 taxa and 4.83 million reads.

#### Tables

Table S1. List of taxonomic assignments not resolved to the species level. The table includes the assigned taxon, the highest level of taxonomic resolution, and the potential species contributing to that assignment. When multiple ASVs matched the same genus or family group, a numeric identifier was added to differentiate the distinct species groups.

| **taxon** | **taxonomic_level** | **species** | **genus** | **family** | **order** |
| --- | --- | --- | --- | --- | --- |
| Anoplarchus | genus | Anoplarchus insignis | Anoplarchus | Stichaeidae | Perciformes |
| Anoplarchus | genus | Anoplarchus purpurescens | Anoplarchus | Stichaeidae | Perciformes |
| Cottus | genus | Cottus asper | Cottus | Cottidae | Perciformes |
| Cottus | genus | Cottus beldingii | Cottus | Cottidae | Perciformes |
| Cottus | genus | Cottus gulosus | Cottus | Cottidae | Perciformes |
| Cottus | genus | Cottus perplexus | Cottus | Cottidae | Perciformes |
| Cottus | genus | Cottus aleuticus | Cottus | Cottidae | Perciformes |
| Cottus | genus | Cottus bairdii | Cottus | Cottidae | Perciformes |
| Cottus | genus | Cottus bendirei | Cottus | Cottidae | Perciformes |
| Cottus | genus | Cottus cognatus | Cottus | Cottidae | Perciformes |
| Cottus | genus | Cottus confusus | Cottus | Cottidae | Perciformes |
| Cottus | genus | Cottus pitensis | Cottus | Cottidae | Perciformes |
| Cottus | genus | Cottus rhotheus | Cottus | Cottidae | Perciformes |
| Cottus_Leptocottus | genus | Cottus asper | Cottus | Cottidae | Perciformes |
| Cottus_Leptocottus | genus | Cottus beldingii | Cottus | Cottidae | Perciformes |
| Cottus_Leptocottus | genus | Cottus gulosus | Cottus | Cottidae | Perciformes |
| Cottus_Leptocottus | genus | Cottus marginatus | Cottus | Cottidae | Perciformes |
| Cottus_Leptocottus | genus | Cottus perplexus | Cottus | Cottidae | Perciformes |
| Cottus_Leptocottus | genus | Cottus ricei | Cottus | Cottidae | Perciformes |
| Cottus_Leptocottus | genus | Leptocottus armatus | Leptocottus | Cottidae | Perciformes |
| Cottus_Leptocottus | genus | Cottus aleuticus | Cottus | Cottidae | Perciformes |
| Cottus_Leptocottus | genus | Cottus bairdii | Cottus | Cottidae | Perciformes |
| Cottus_Leptocottus | genus | Cottus bendirei | Cottus | Cottidae | Perciformes |
| Cottus_Leptocottus | genus | Cottus cognatus | Cottus | Cottidae | Perciformes |
| Cottus_Leptocottus | genus | Cottus confusus | Cottus | Cottidae | Perciformes |
| Cottus_Leptocottus | genus | Cottus pitensis | Cottus | Cottidae | Perciformes |
| Cottus_Leptocottus | genus | Cottus rhotheus | Cottus | Cottidae | Perciformes |
| Cottus_Leptocottus | genus | Cottus carolinae | Cottus | Cottidae | Perciformes |
| Cottus_Leptocottus | genus | Cottus specus | Cottus | Cottidae | Perciformes |
| Gadus | genus | Gadus chalcogrammus | Gadus | Gadidae | Gadiformes |
| Gadus | genus | Gadus macrocephalus | Gadus | Gadidae | Gadiformes |
| Hexagrammos | genus | Hexagrammos lagocephalus | Hexagrammos | Hexagrammidae | Perciformes |
| Hexagrammos | genus | Hexagrammos octogrammus | Hexagrammos | Hexagrammidae | Perciformes |
| Hexagrammos | genus | Hexagrammos stelleri | Hexagrammos | Hexagrammidae | Perciformes |
| Hypomesus | genus | Hypomesus pretiosus | Hypomesus | Osmeridae | Osmeriformes |
| Hypomesus | genus | Hypomesus transpacificus | Hypomesus | Osmeridae | Osmeriformes |
| Lepidopsetta | genus | Lepidopsetta bilineata | Lepidopsetta | Pleuronectidae | Pleuronectiformes |
| Lepidopsetta | genus | Lepidopsetta polyxystra | Lepidopsetta | Pleuronectidae | Pleuronectiformes |
| Myoxocephalus | genus | Myoxocephalus jaok | Myoxocephalus | Cottidae | Perciformes |
| Myoxocephalus | genus | Myoxocephalus polyacanthocephalus | Myoxocephalus | Cottidae | Perciformes |
| Myoxocephalus | genus | Myoxocephalus stelleri | Myoxocephalus | Cottidae | Perciformes |
| Myoxocephalus | genus | Myoxocephalus quadricornis | Myoxocephalus | Cottidae | Perciformes |
| Myoxocephalus | genus | Myoxocephalus scorpius | Myoxocephalus | Cottidae | Perciformes |
| Myoxocephalus | genus | Myoxocephalus thompsonii | Myoxocephalus | Cottidae | Perciformes |
| Oncorhynchus | genus | Oncorhynchus mykiss | Oncorhynchus | Salmonidae | Salmoniformes |
| Oncorhynchus | genus | Oncorhynchus nerka | Oncorhynchus | Salmonidae | Salmoniformes |
| Oncorhynchus | genus | Oncorhynchus keta | Oncorhynchus | Salmonidae | Salmoniformes |
| Oncorhynchus | genus | Oncorhynchus gorbuscha | Oncorhynchus | Salmonidae | Salmoniformes |
| Oncorhynchus | genus | Oncorhynchus kisutch | Oncorhynchus | Salmonidae | Salmoniformes |
| Pleuronectidae 1 | family | Hippoglossoides elassodon | Hippoglossoides | Pleuronectidae | Pleuronectiformes |
| Pleuronectidae 1 | family | Hippoglossoides robustus | Hippoglossoides | Pleuronectidae | Pleuronectiformes |
| Pleuronectidae 1 | family | Limanda aspera | Limanda | Pleuronectidae | Pleuronectiformes |
| Pleuronectidae 2 | family | Isopsetta isolepis | Isopsetta | Pleuronectidae | Pleuronectiformes |
| Pleuronectidae 2 | family | Parophrys vetulus | Parophrys | Pleuronectidae | Pleuronectiformes |
| Pleuronectidae 2 | family | Psettichthys melanostictus | Psettichthys | Pleuronectidae | Pleuronectiformes |
| Pleuronectidae 3 | family | Liopsetta glacialis | Liopsetta | Pleuronectidae | Pleuronectiformes |
| Pleuronectidae 3 | family | Platichthys stellatus | Platichthys | Pleuronectidae | Pleuronectiformes |
| Salvelinus | genus | Salvelinus alpinus | Salvelinus | Salmonidae | Salmoniformes |
| Salvelinus | genus | Salvelinus malma | Salvelinus | Salmonidae | Salmoniformes |
| Salvelinus | genus | Salvelinus confluentus | Salvelinus | Salmonidae | Salmoniformes |
| Sebastes 1 | genus | Sebastes caurinus | Sebastes | Sebastidae | Perciformes |
| Sebastes 1 | genus | Sebastes maliger | Sebastes | Sebastidae | Perciformes |
| Sebastes 2 | genus | Sebastes auriculatus | Sebastes | Sebastidae | Perciformes |
| Sebastes 2 | genus | Sebastes rastrelliger | Sebastes | Sebastidae | Perciformes |
| Sebastes 3 | genus | Sebastes aleutianus | Sebastes | Sebastidae | Perciformes |
| Sebastes 3 | genus | Sebastes alutus | Sebastes | Sebastidae | Perciformes |
| Sebastes 3 | genus | Sebastes babcocki | Sebastes | Sebastidae | Perciformes |
| Sebastes 3 | genus | Sebastes brevispinis | Sebastes | Sebastidae | Perciformes |
| Sebastes 3 | genus | Sebastes elongatus | Sebastes | Sebastidae | Perciformes |
| Sebastes 3 | genus | Sebastes emphaeus | Sebastes | Sebastidae | Perciformes |
| Sebastes 3 | genus | Sebastes flavidus | Sebastes | Sebastidae | Perciformes |
| Sebastes 3 | genus | Sebastes goodei | Sebastes | Sebastidae | Perciformes |
| Sebastes 3 | genus | Sebastes helvomaculatus | Sebastes | Sebastidae | Perciformes |
| Sebastes 3 | genus | Sebastes melanops | Sebastes | Sebastidae | Perciformes |
| Sebastes 3 | genus | Sebastes melanostictus | Sebastes | Sebastidae | Perciformes |
| Sebastes 3 | genus | Sebastes miniatus | Sebastes | Sebastidae | Perciformes |
| Sebastes 3 | genus | Sebastes pinniger | Sebastes | Sebastidae | Perciformes |
| Sebastes 3 | genus | Sebastes proriger | Sebastes | Sebastidae | Perciformes |
| Sebastes 3 | genus | Sebastes rosaceus | Sebastes | Sebastidae | Perciformes |
| Sebastes 3 | genus | Sebastes ruberrimus | Sebastes | Sebastidae | Perciformes |
| Sebastes 3 | genus | Sebastes variegatus | Sebastes | Sebastidae | Perciformes |
| Sebastes 3 | genus | Sebastes wilsoni | Sebastes | Sebastidae | Perciformes |
| Sebastes 3 | genus | Sebastes zacentrus | Sebastes | Sebastidae | Perciformes |
| Sebastes 3 | genus | Sebastes nigrocinctus | Sebastes | Sebastidae | Perciformes |
| Sebastes 3 | genus | Sebastes melanostomus | Sebastes | Sebastidae | Perciformes |
| Sebastes 3 | genus | Sebastes paucispinis | Sebastes | Sebastidae | Perciformes |
| Sebastolobus | genus | Sebastolobus alascanus | Sebastolobus | Sebastidae | Perciformes |
| Sebastolobus | genus | Sebastolobus altivelis | Sebastolobus | Sebastidae | Perciformes |
| Sebastolobus | genus | Sebastolobus macrochir | Sebastolobus | Sebastidae | Perciformes |
| Stichaeidae | family | Leptoclinus maculatus | Leptoclinus | Stichaeidae | Perciformes |
| Stichaeidae | family | Lumpenus fabricii | Lumpenus | Stichaeidae | Perciformes |
| Xiphister | genus | Xiphister atropurpureus | Xiphister | Stichaeidae | Perciformes |
| Xiphister | genus | Xiphister mucosus | Xiphister | Stichaeidae | Perciformes |

Table S2. Species compositions of the 13 mock communities created to evaluate amplification efficiency. Values represent the proportion (%) of each of the 22 species included in the mock community.

| **Scientific name** | **Common name** | **COM1** | **COM2** | **COM3** | **COM4** | **COM5** | **COM6** | **COM7** | **COM8** | **COM9** | **COM10** | **COM11** | **COM12** | **COM13** |
| --- | --- | --- | --- | --- | --- | --- | --- | --- | --- | --- | --- | --- | --- | --- |
| *Ammodytes personatus* | Pacific sandlance | 9.7 | 28.7 | 0 | 0 | 0 | 0 | 10.6 | 6.7 | 20 | 0 | 9.2 | 10.2 | 0 |
| *Blepsias cirrhosus* | Silverspotted sculpin | 10.5 | 0 | 0 | 0 | 5.5 | 0 | 1.1 | 3.6 | 4.3 | 0 | 0 | 5.5 | 4.8 |
| *Clupea pallasii* | Pacific herring | 0 | 4.6 | 33.5 | 0 | 0 | 0 | 0 | 5.3 | 0 | 0 | 0 | 0 | 0 |
| *Cymatogaster aggregata* | Shiner perch | 9.7 | 0 | 0 | 0 | 25.6 | 56.7 | 0 | 16.7 | 20 | 0 | 9.2 | 0 | 8.9 |
| *Gadus macrocephalus* | Pacific cod | 0 | 0 | 0 | 7.6 | 0 | 0 | 3.7 | 6.3 | 7.6 | 10.4 | 0 | 0 | 0 |
| *Gasterosteus aculeatus* | Three-spine stickleback | 0 | 0 | 0 | 0 | 0 | 0 | 0 | 0 | 0 | 0 | 44.4 | 24.5 | 64.5 |
| *Hexagrammos stelleri* | Whitespotted greenling | 10 | 0 | 4.3 | 0 | 5.2 | 4 | 3.9 | 6.8 | 8.2 | 11.2 | 9.4 | 10.4 | 4.6 |
| *Lepidopsetta bilineata* | Pacific rock sole | 0 | 2.8 | 4.1 | 0 | 0 | 2.5 | 0 | 3.2 | 0 | 10.6 | 0 | 0 | 0 |
| *Leptocottus armatus* | Pacific staghorn sculpin | 9.2 | 5.4 | 4 | 0 | 12.1 | 18.5 | 0 | 0 | 0 | 0 | 4.4 | 0 | 0 |
| *Limanda aspera* | Yellowfin sole | 0 | 2.7 | 4 | 3.8 | 2.4 | 1.1 | 0 | 0 | 0 | 0 | 0 | 0 | 4.3 |
| *Lumpenus sagitta* | Snake prickleback | 11.3 | 0 | 9.8 | 0 | 14.9 | 12 | 2.5 | 7.8 | 0 | 0 | 10.7 | 0 | 0 |
| *Oncorhynchus gorbuscha* | Pink salmon | 0 | 20.8 | 15.3 | 28.9 | 0 | 0 | 0.8 | 12.1 | 2.9 | 0 | 3.3 | 3.7 | 0 |
| *Oncorhynchus keta* | Chum salmon | 10.2 | 29.9 | 8.8 | 20.8 | 0 | 0 | 0 | 7 | 0 | 0 | 0 | 0 | 0 |
| *Oncorhynchus kisutch* | Coho salmon | 9 | 0 | 3.9 | 18.4 | 4.7 | 0 | 0 | 0 | 0 | 0 | 0 | 0 | 4.1 |
| *Oncorhynchus nerka* | Sockeye salmon | 0 | 0 | 0 | 9.8 | 0 | 0 | 0 | 0 | 0 | 0 | 0 | 0 | 0 |
| *Oncorhynchus tshawytscha* | Chinook salmon | 0 | 0 | 0 | 6.5 | 0 | 0 | 0 | 0 | 0 | 0 | 0 | 0 | 0 |
| *Ophiodon elongatus* | Lingcod | 0 | 0 | 0 | 0 | 4.3 | 0 | 41.9 | 5.6 | 16.9 | 9.3 | 0 | 4.3 | 0 |
| *Pholis laeta* | Crescent gunnels | 10.5 | 0 | 0 | 4.3 | 0 | 3.3 | 31.6 | 3.6 | 8.6 | 0 | 0 | 11 | 0 |
| *Sebastes auriculatus* | Brown rockfish | 0 | 3 | 0 | 0 | 13.3 | 0 | 0 | 0 | 0 | 22.8 | 0 | 0 | 4.6 |
| *Sebastes caurinus* | Copper rockfish | 0 | 0 | 3.9 | 0 | 11.8 | 0 | 1.6 | 6.2 | 7.4 | 20.3 | 0 | 4.7 | 4.1 |
| *Sebastes flavidus* | Yellowtail rockfish | 0 | 2 | 0 | 0 | 0 | 0 | 0 | 2.4 | 0 | 15.5 | 0 | 0 | 0 |
| *Syngnathus leptorhynchus* | Bay pipefish | 9.8 | 0 | 8.5 | 0 | 0 | 1.9 | 2.1 | 6.7 | 4 | 0 | 9.3 | 25.6 | 0 |

Table S3. Species functional traits for each trait category (schooling behavior, position-in-water-column, and scale-type) for species either identified by eDNA or beach seine methods.

| **Taxonomic group** | **Scientific name** | **Common name** | **Schooling behavior** | **Position-in-water-column** | **Scale type** |
| --- | --- | --- | --- | --- | --- |
| *Clupea pallasii* |  | Pacific herring | obligatory schooling | pelagic | deciduous scales |
| *Trichodon trichodon* |  | Pacific sandfish | obligatory schooling | both | no scales |
| *Clinocottus acuticeps* |  | Sharpnose sculpin | solitary | demersal | no scales |
| *Clinocottus embryum* |  | Calico sculpin | solitary | demersal | no scales |
| *Cottus* | *Cottus asper*  *Cottus beldingii*  *Cottus gulosus*  *Cottus perplexus*  *Cottus aleuticus*  *Cottus bairdii*  *Cottus bendirei*  *Cottus cognatus*  *Cottus confusus*  *Cottus pitensis*  *Cottus rhotheus* | Sculpins | solitary | demersal | no scales |
| *Cottus_Leptocottus* | *Cottus asper*  *Cottus beldingii*  *Cottus gulosus*  *Cottus marginatus*  *Cottus perplexus*  *Cottus ricei*  *Leptocottus armatus*  *Cottus aleuticus*  *Cottus bairdii*  *Cottus bendirei*  *Cottus cognatus*  *Cottus confusus*  *Cottus pitensis*  *Cottus rhotheus*  *Cottus carolinae*  *Cottus specus* | Sculpins | solitary | demersal | no scales |
| *Gobiesox maeandricus* |  | Northern clingfish | solitary | demersal | no scales |
| *Liparis mucosus* |  | Slimy snailfish | solitary | demersal | no scales |
| *Myoxocephalus* | *Myoxocephalus jaok*  *Myoxocephalus polyacanthocephalus*  *Myoxocephalus stelleri*  *Myoxocephalus quadricornis*  *Myoxocephalus scorpius*  *Myoxocephalus thompsonii* | Sculpins | solitary | demersal | no scales |
| *Oligocottus maculosus* |  | Tidepool sculpin | solitary | demersal | no scales |
| *Oligocottus snyderi* |  | Fluffy sculpin | solitary | demersal | no scales |
| *Gasterosteus aculeatus* |  | Three-spine stickleback | obligatory schooling | both | plates |
| *Agonopsis vulsa* |  | Northern spearnose poacher | solitary | demersal | plates |
| *Blepsias bilobus* |  | Crested sculpin | solitary | demersal | plates |
| *Blepsias cirrhosus* |  | Silverspotted sculpin | solitary | demersal | plates |
| *Nautichthys oculofasciatus* |  | Sailfin sculpin | solitary | demersal | plates |
| *Pallasina barbata* |  | Tubenose poacher | solitary | demersal | plates |
| *Podothecus accipenserinus* |  | Sturgeon poacher | solitary | demersal | plates |
| *Rhamphocottus richardsonii* |  | Grunt sculpin | solitary | demersal | plates |
| *Aulorhynchus flavidus* |  | Tubesnout | obligatory schooling | pelagic | plates |
| *Syngnathus leptorhynchus* |  | Bay pipefish | solitary | pelagic | plates |
| *Ammodytes personatus* |  | Pacific sandlance | obligatory schooling | both | scales |
| *Anoplarchus* | *Anoplarchus insignis*  *Anoplarchus purpurescens* | Slender coxcomb or High coxcomb | solitary | demersal | scales |
| *Anoplopoma fimbria* |  | Sablefish | solitary | demersal | scales |
| *Apodicthys flavidus* |  | Penpoint gunnel | solitary | demersal | scales |
| *Artedius spp.* | *Artedius fenestralis*  *Artedius harringtoni*  *Artedius lateralis* | Artedius sculpins | solitary | demersal | scales |
| *Chitonotus pugetensis* |  | Roughback sculpin | solitary | demersal | scales |
| *Citharichthys sordidus* |  | Pacific sanddab | solitary | demersal | scales |
| *Citharichthys stigmaeus* |  | Speckled sanddab | solitary | demersal | scales |
| *Gadus* | *Gadus chalcogrammus, Gadus macrocephalus* | Cods | solitary | demersal | scales |
| *Hemilepidotus spp.* | *Hemilepidotus hemilepidotus* | Irish lords | solitary | demersal | scales |
| *Hexagrammos* | *Hexagrammos lagocephalus, Hexagrammos octogrammus, Hexagrammos stelleri* | Greenlings | solitary | demersal | scales |
| *Hexagrammos decagrammus* |  | Kelp greenling | solitary | demersal | scales |
| *Hippoglossus stenolepis* |  | Pacific halibut | solitary | demersal | scales |
| *Lepidogobius lepidus* |  | Bay goby | solitary | demersal | scales |
| *Lepidopsetta* | *Lepidopsetta bilineata*  *Lepidopsetta polyxystra* | Righteye flounders | solitary | demersal | scales |
| *Lumpenus sagitta* |  | Snake prickleback | solitary | demersal | scales |
| *Microgadus proximus* |  | Pacific tomcod | solitary | demersal | scales |
| *Ophiodon elongatus* |  | Lingcod | solitary | demersal | scales |
| *Oxylebius pictus* |  | Painted greenling | solitary | demersal | scales |
| *Pholis laeta* |  | Crescent gunnel | solitary | demersal | scales |
| *Phytichthys chirus* |  | Ribbon prickleback | solitary | demersal | scales |
| *Pleuronectidae 1* | *Hippoglossoides elassodon*  *Hippoglossoides robustus*  *Limanda aspera* |  | solitary | demersal | scales |
| *Pleuronectidae 2* | *Isopsetta isolepis*  *Parophrys vetulus*  *Psettichthys melanostictus* |  | solitary | demersal | scales |
| *Pleuronectidae 3* | *Liopsetta glacialis*  *Platichthys stellatus* |  | solitary | demersal | scales |
| *Pleuronichthys coenosus* |  | C-O sole | solitary | demersal | scales |
| *Rhinogobiops nicholsii* |  | Blackeye goby | solitary | demersal | scales |
| *Ronquilus jordani* |  | Northern ronquil | solitary | demersal | scales |
| *Sebastes 1* | *Sebastes caurinus*  *Sebastes maliger* | Copper or Quillback rockfish | solitary | demersal | scales |
| *Sebastes 2* | *Sebastes auriculatus*  *Sebastes rastrelliger* | Brown rockfish Grass rockfish | solitary | demersal | scales |
| *Sebastolobus* | *Sebastolobus alascanus*  *Sebastolobus altivelis*  *Sebastolobus macrochir* | Thornyheads | solitary | demersal | scales |
| *Stichaeidae* | *Leptoclinus maculatus*  *Lumpenus fabricii* | Daubed shanny or Slender eelblenny | solitary | demersal | scales |
| *Stichaeus punctatus* |  | Arctic shanny | solitary | demersal | scales |
| *Xiphister* | *Xiphister atropurpureus*  *Xiphister mucosus* | Black prickleback or Rock prickleback | solitary | demersal | scales |
| *Sebastes 3* | *Sebastes aleutianus*  *Sebastes alutus*  *Sebastes babcocki*  *Sebastes brevispinis*  *Sebastes elongatus*  *Sebastes emphaeus*  *Sebastes flavidus*  *Sebastes goodei*  *Sebastes helvomaculatus*  *Sebastes melanops*  *Sebastes melanostictus*  *Sebastes miniatus*  *Sebastes pinniger*  *Sebastes proriger*  *Sebastes rosaceus*  *Sebastes ruberrimus*  *Sebastes variegatus*  *Sebastes wilsoni*  *Sebastes zacentrus*  *Sebastes nigrocinctus*  *Sebastes melanostomus*  *Sebastes paucispinis*  *Sebastes entomelas*  *Sebastes nigrocinctus* | Pacific ocean perch,  Redbanded rockfish,  Greenstriped rockfish,  Yellowtail rockfish,  Black rockfish,  Blackspotted rockfish,  Boccaio, or  Redstripe rockfish  Yelloweye rockfish  Pygmy rockfish | obligatory schooling | pelagic | scales |
| *Brachyistius frenatus* |  | Kelp perch | faculative schooling | pelagic | scales |
| *Cymatogaster aggregata* |  | Shiner perch | faculative schooling | pelagic | scales |
| *Embiotoca lateralis* |  | Striped surfperch | faculative schooling | pelagic | scales |
| *Hypomesus* | *Hypomesus pretiosus*  *Hypomesus transpacificus* | Smelts | obligatory schooling | pelagic | scales |
| *Mallotus villosus* |  | Capelin | obligatory schooling | pelagic | scales |
| *Oncorhynchus* | *Oncorhynchus mykiss*  *Oncorhynchus nerka*  *Oncorhynchus keta*  *Oncorhynchus gorbuscha*  *Oncorhynchus kisutch* | Salmon and Trouts | obligatory schooling | pelagic | scales |
| *Oncorhynchus clarkii* |  | Cutthroat trout | obligatory schooling | pelagic | scales |
| *Oncorhynchus tshawytscha* |  | King salmon | obligatory schooling | pelagic | scales |
| *Salvelinus* | *Salvelinus alpinus*  *Salvelinus malma*  *Salvelinus confluentus* | Chars | obligatory schooling | pelagic | scales |
| *Zaprora silenus* |  | Prowfish | solitary | pelagic | scales |
| *Cryptacanthodes giganteus* |  | Giant wrymouth | solitary | demersal | some scales |
| *Enophrys bison* |  | Buffalo sculpin | solitary | demersal | some scales |
| *Enophrys lucasi* |  | Leister sculpin | solitary | demersal | some scales |
| *Scorpaenichthys marmoratus* |  | Cabezon | solitary | demersal | some scales |
| *Synchirus gilli* |  | Manacled sculpin | solitary | demersal | some scales |

Table S4. Permutation-based ANOVA (PERMANOVA) of logical pairwise comparisons (contrasts) of the interaction between groups of habitat and gear type for fish community composition.

| Contrasts (logical comparisons) | DF | Marginal R^2^ | Sum of squares | Psuedo-F statistic | P-value |
| --- | --- | --- | --- | --- | --- |
| **Eelgrass eDNA and seine** | 1 | 0.075 | 1.622 | 5.222 | 0.0005 |
| residual | 64 | 0.925 | 19.875 |  |  |
| total | 65 | 1 | 21.496 |  |  |
| **Kelp eDNA and seine** | 1 | 0.113 | 1.08 | 3.302 | 0.0005 |
| residual | 26 | 0.887 | 8.503 |  |  |
| total | 27 | 1 | 9.583 |  |  |
| **Mixed eelgrass eDNA and seine** | 1 | 0.127 | 1.634 | 5.977 | 0.0005 |
| residual | 41 | 0.873 | 11.209 |  |  |
| total | 42 | 1 | 12.843 |  |  |
| **eDNA: eelgrass and kelp** | 1 | 0.057 | 1.312 | 4.180 | 0.0005 |
| residual | 68 | 0.942 | 21.340 |  |  |
| total | 69 | 1 | 22.652 |  |  |
| **eDNA: eelgrass and mixed eelgrass** | 1 | 0.33 | 0.839 | 2.682 | 0.0005 |
| residual | 79 | 0.967 | 24.696 |  |  |
| total | 80 | 1 | 25.535 |  |  |
| **eDNA: kelp and mixed eelgrass** | 1 | 0.068 | 1.150 | 3.750 | 0.0005 |
| residual | 51 | 0.932 | 15.636 |  |  |
| total | 52 | 1 | 16.786 |  |  |
| **Seine: eelgrass and kelp** | 1 | 0.103 | 0.81 | 2.532 | 0.004 |
| residual | 22 | 0.897 | 7.038 |  |  |
| total | 23 | 1 | 7.848 |  |  |
| **Seine: eelgrass and mixed eelgrass** | 1 | 0.096 | 0.696 | 2.652 | 0.0035 |
| residual | 26 | 0.907 | 6.822 |  |  |
| total | 27 | 1 | 7.518 |  |  |
| **Seine: kelp and mixed eelgrass** | 1 | 0.2 | 1.021 | 4.007 | 0.0005 |
| residual | 16 | 0.799 | 4.075 |  |  |
| total | 17 | 1 | 5.096 |  |  |

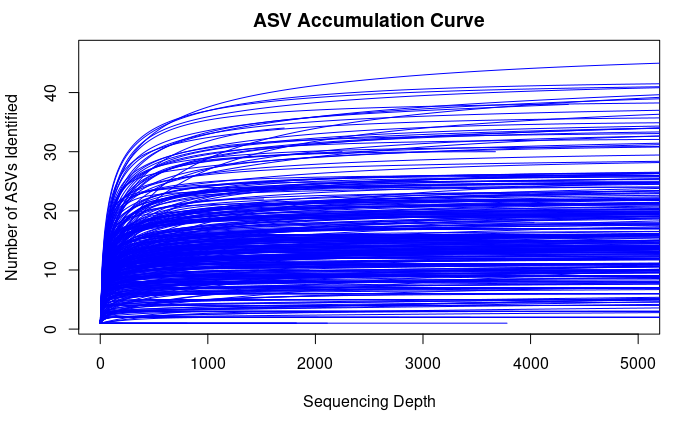

#### Figures

Figure S1. Accumulation curve of the relationship between sequencing depth and the number of amplicon sequence variants (ASV) identified using the R package *vegan*.

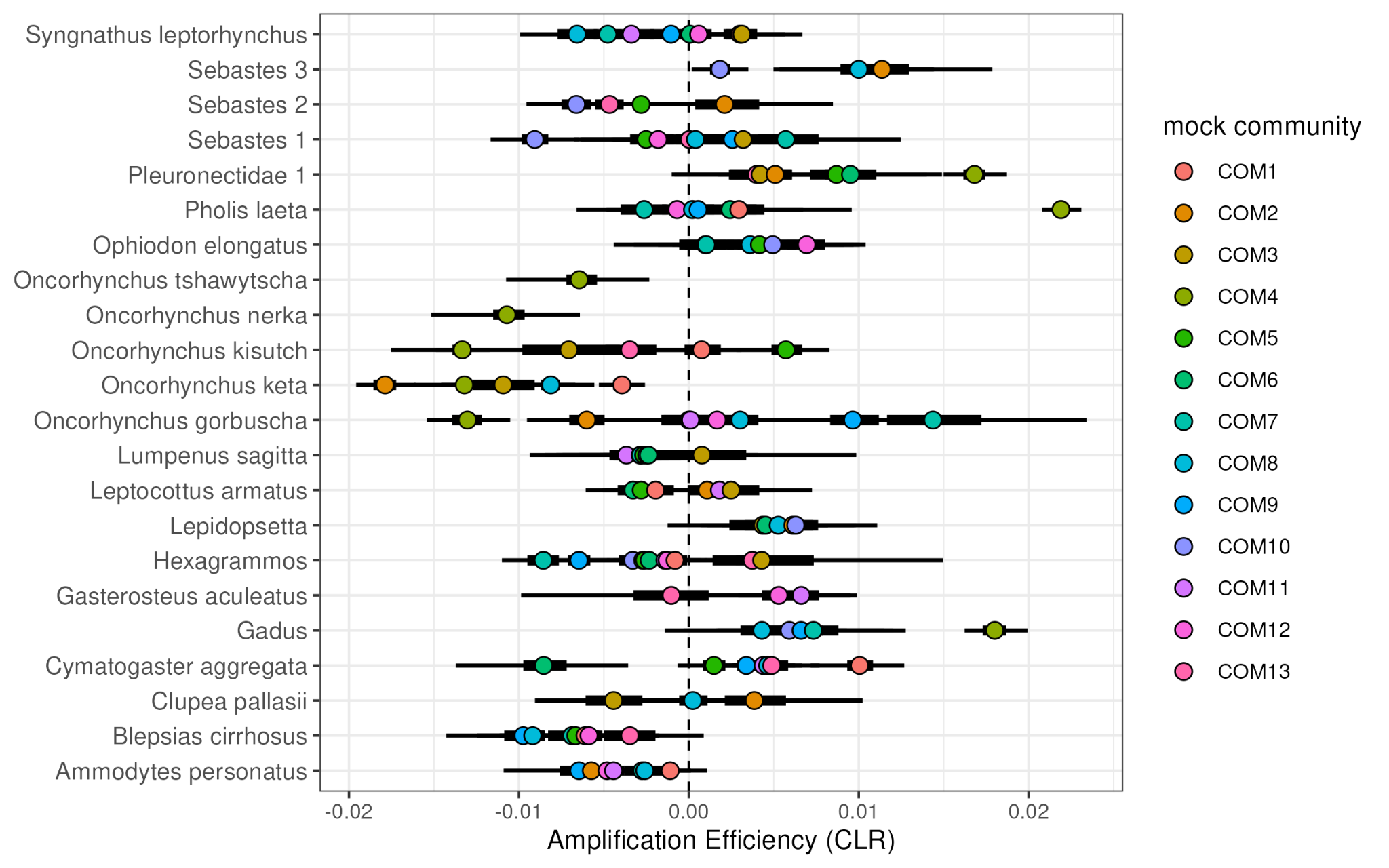
Figure S2. Centered log-ratio (CLR) transformation of amplification efficiencies estimates for 22 taxa from each of the 13 mock communities.

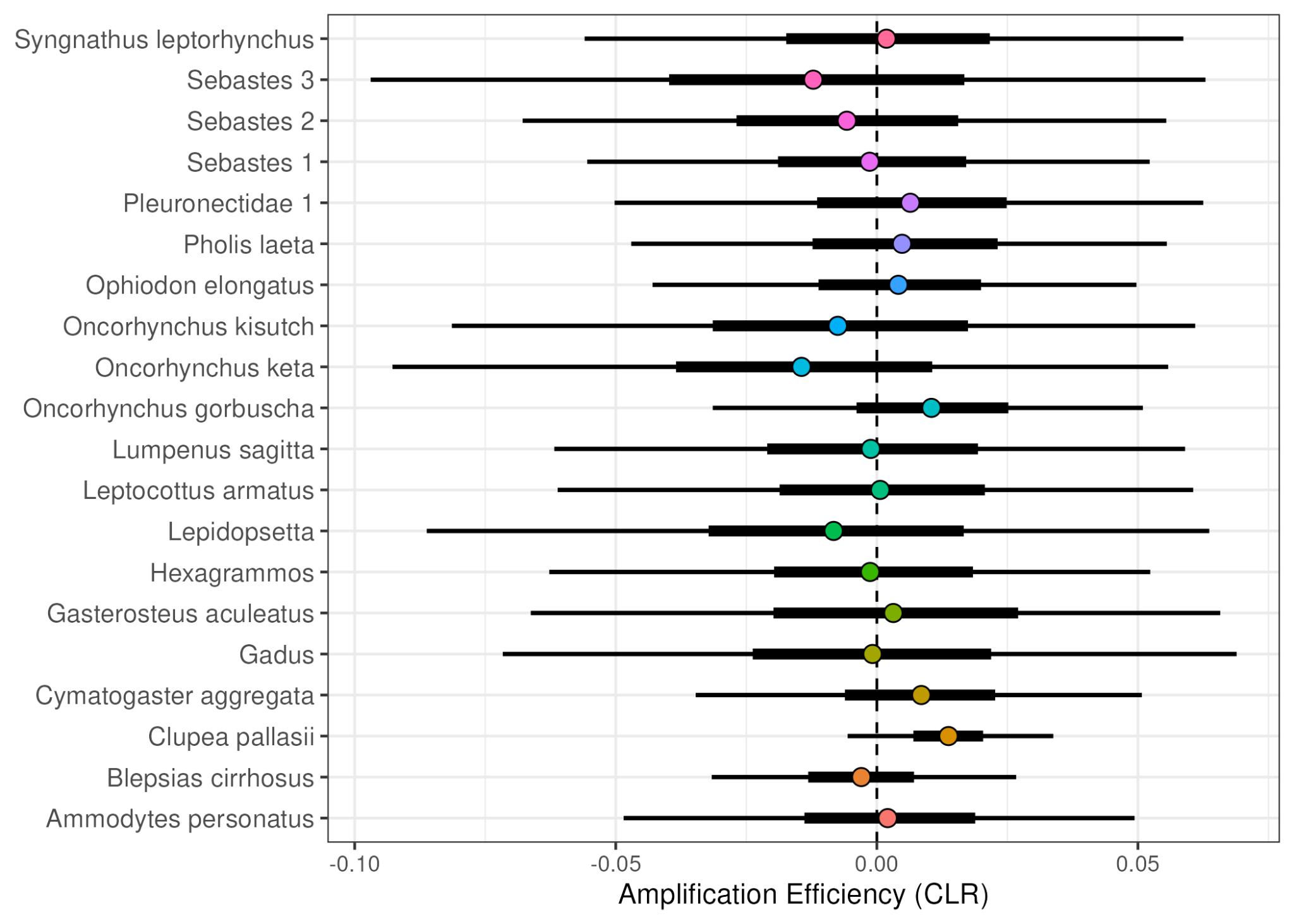

Figure S3. Comparison of relative amplification efficiencies across 20 taxa in the mock communities. Interquartile ranges (IQRs) of amplification efficiency estimates for each taxon derived from 12 mock communities overlap both each other as well as zero, indicating no substation variation among taxa. Points represent posterior mean estimates of relative amplification efficiency after centered log-ratio transformation (CLR). Thick horizontal lines represent 50% IQRs, while thin lines represent 95% IQRs. The dashed vertical line indicates the mean amplification efficiency across all species.

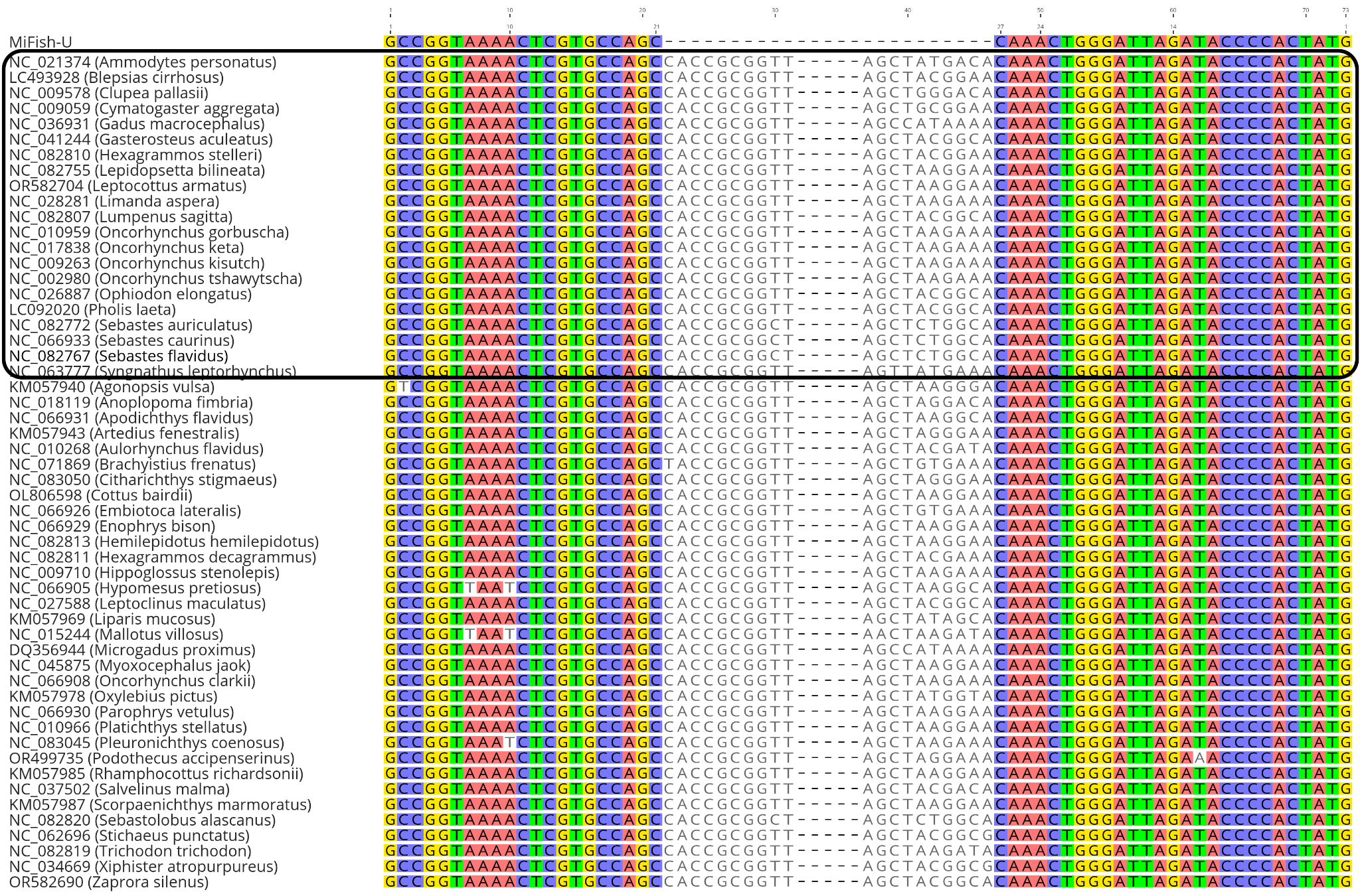

Figure S4. Alignment of 12S MiFish-U primer-binding regions for species with available 12S sequences spanning the binding sites. Species included in mock communities are highlighted in the black box. Colored bases indicate the matching base pair positions between the primer and species sequences. Accession numbers for the reference sequences are provided alongside the species names.

Figure S5. Species accumulation curves in eelgrass (left, green), understory kelp (middle, orange), and mixed eelgrass meadows (right, yellow) for sites sampled with eDNA (solid lines, light ribbon) and beach seines (dashed lines, dark ribbon).
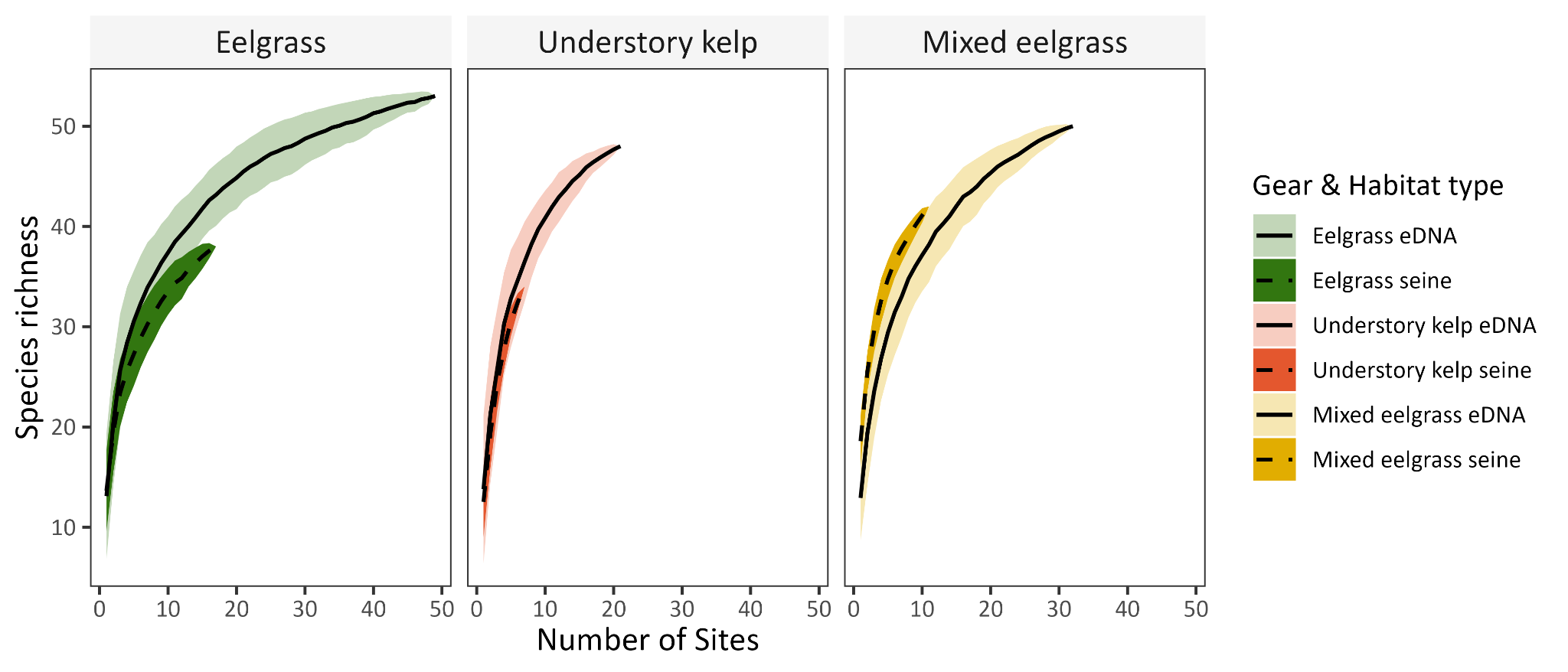

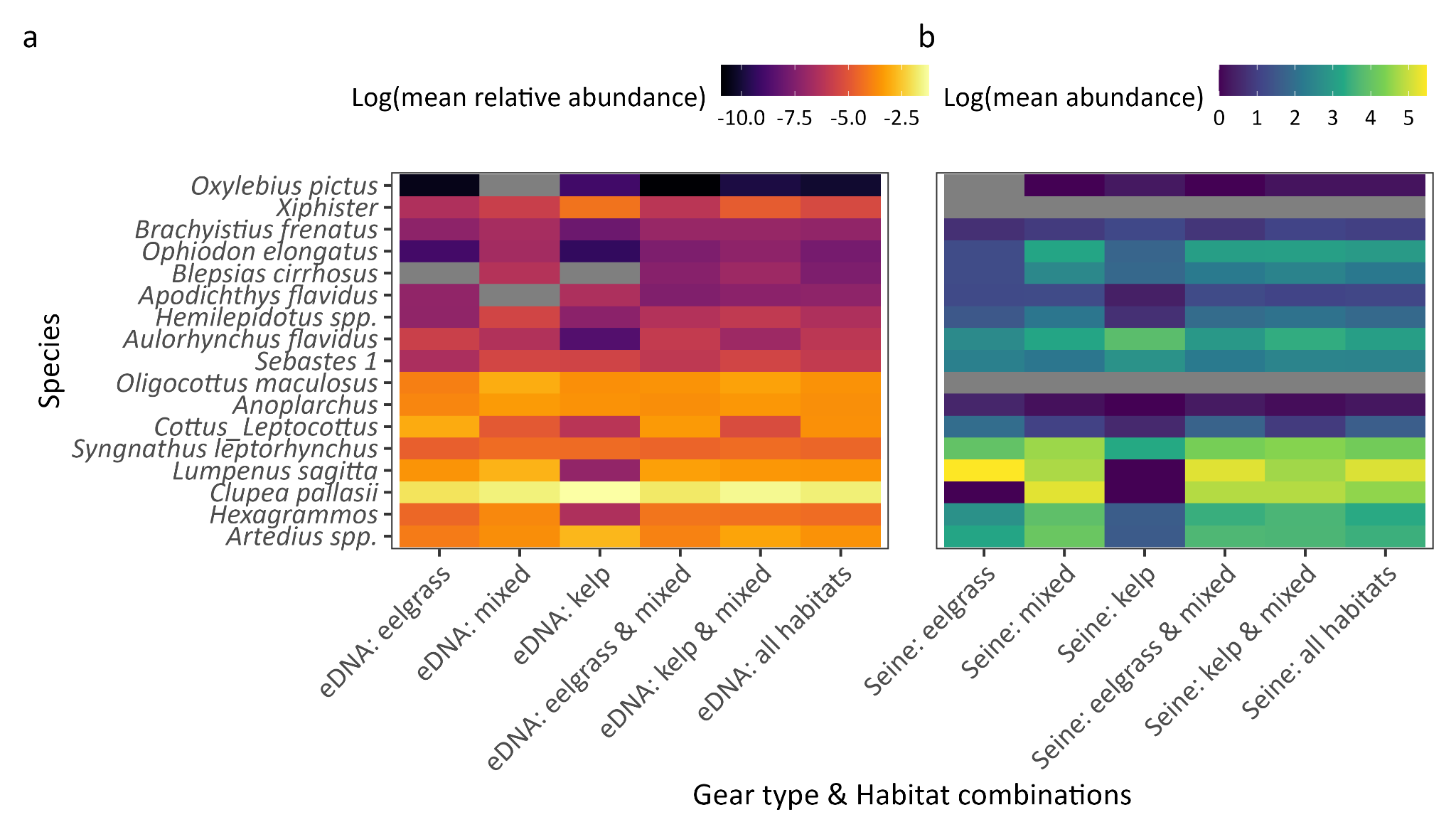
Figure S6. Logarithmic-transformed mean relative abundance of indicator species sampled with eDNA (a) and logarithmic-transformed mean absolute abundance of indicator species sampled with beach seines (b) in gear and habitat combinations. Gray indicates species were absent for said gear and habitat combination. Cool colors represent low mean relative/absolute abundance and warm colors represent high mean relative/absolute abundance.

### Supplementary Code

\# PPC function with separate p-values for each data source

ppcOcc_intMsPGOcc <- function(object, fit.stat = "chi-squared", thin.by = 1) {

\# Input validation

if (missing(object)) {

stop("error: object must be specified")

}

if (class(object) != "intMsPGOcc") {

stop("error: object must be of class 'intMsPGOcc'")

}

if (!tolower(fit.stat) %in% c("chi-squared", "freeman-tukey", "chi-square")) {

stop("error: fit.stat must be either 'chi-squared' or 'freeman-tukey'")

}

fit.stat <- tolower(fit.stat)

\# Helper functions

logit_inv <- function(z, a = 0, b = 1) {

b - (b - a) / (1 + exp(z))

}

cat("Running posterior predictive check for intMsPGOcc output...\n")

\# Extract model components

y <- object$y

sites <- object$sites

species_list <- object$species

X_p <- object$X.p

n_data <- length(y)

n_samples_total <- object$n.post * object$n.chains

\# Apply thinning

sample_indices <- seq(1, n_samples_total, by = thin.by)

n_samples <- length(sample_indices)

cat("Using", n_samples, "samples (every", thin.by, "of", n_samples_total, "total samples)\n")

cat("Calculating p-values for", n_data, "data sources\n")

\# Extract thinned samples

z_samples <- object$z.samples[sample_indices, , , drop = FALSE]

alpha_samples <- object$alpha.samples[sample_indices, , drop = FALSE]

\# Initialize storage - separate for each data source

chi_sq_obs_by_source <- matrix(0, nrow = n_samples, ncol = n_data)

chi_sq_rep_by_source <- matrix(0, nrow = n_samples, ncol = n_data)

chi_sq_obs_total <- numeric(n_samples)

chi_sq_rep_total <- numeric(n_samples)

epsilon <- 1e-04

\# Process each MCMC sample

for(s in 1:n_samples) {

if(s %% 50 == 0) cat("Processing sample", s, "of", n_samples, "\n")

sample_chi_obs_total <- 0

sample_chi_rep_total <- 0

\# Process each data source

for(d in 1:n_data) {

sample_chi_obs_d <- 0 # Chi-square for this data source

sample_chi_rep_d <- 0 # Chi-square replicated for this data source

y_curr <- y[[d]]

sites_curr <- sites[[d]]

species_curr <- species_list[[d]]

X_p_curr <- X_p[[d]]

\# Map species to full community indices

if(is.character(species_curr)) {

sp_indices <- match(species_curr, object$sp.names)

} else {

sp_indices <- species_curr

}

n_species_d <- length(sp_indices)

\# Get alpha coefficients for this data source

alpha_start_idx <- 1

if(d > 1) {

for(prev_d in 1:(d-1)) {

alpha_start_idx <- alpha_start_idx +

length(species_list[[prev_d]]) * ncol(X_p[[prev_d]])

}

}

n_alpha_curr <- n_species_d * ncol(X_p_curr)

alpha_curr <- alpha_samples[s, alpha_start_idx:(alpha_start_idx + n_alpha_curr - 1)]

alpha_matrix <- matrix(alpha_curr, nrow = n_species_d, ncol = ncol(X_p_curr), byrow = TRUE)

\# Process each species in this data source

for(i in 1:n_species_d) {

sp_idx <- sp_indices[i]

alpha_sp <- alpha_matrix[i, ]

\# Collect all valid observations and expected values for this species

obs_vals <- c()

exp_vals <- c()

rep_vals <- c()

site_rep_ids <- c() # Track which site/rep combination each value comes from

\# Loop through sites and replicates

for(j in 1:length(sites_curr)) {

site_idx <- sites_curr[j]

z_val <- z_samples[s, sp_idx, site_idx]

for(k in 1:dim(y_curr)[3]) {

if(!is.na(y_curr[i, j, k])) {

\# Observed value

obs_vals <- c(obs_vals, y_curr[i, j, k])

\# Expected value

obs_row_idx <- (j - 1) * dim(y_curr)[3] + k

if(obs_row_idx <= nrow(X_p_curr)) {

logit_p <- sum(X_p_curr[obs_row_idx, ] * alpha_sp)

p_detect <- logit_inv(logit_p) * z_val

} else {

next

}

exp_vals <- c(exp_vals, p_detect)

\# Replicated value

rep_vals <- c(rep_vals, rbinom(1, 1, p_detect))

site_rep_ids <- c(site_rep_ids, k) # Group by replicate

}

}

}

\# Group the values and calculate chi-square contributions

if(length(obs_vals) > 0) {

unique_groups <- unique(site_rep_ids)

for(group_id in unique_groups) {

group_mask <- site_rep_ids == group_id

obs_sum <- sum(obs_vals[group_mask])

exp_sum <- sum(exp_vals[group_mask])

rep_sum <- sum(rep_vals[group_mask])

if(exp_sum > 0) {

if(fit.stat %in% c("chi-squared", "chi-square")) {

contrib_obs <- (obs_sum - exp_sum)^2 / (exp_sum + epsilon)

contrib_rep <- (rep_sum - exp_sum)^2 / (exp_sum + epsilon)

} else if(fit.stat == "freeman-tukey") {

contrib_obs <- (sqrt(obs_sum) - sqrt(exp_sum))^2

contrib_rep <- (sqrt(rep_sum) - sqrt(exp_sum))^2

}

\# Add to data source specific totals

sample_chi_obs_d <- sample_chi_obs_d + contrib_obs

sample_chi_rep_d <- sample_chi_rep_d + contrib_rep

}

}

}

}

\# Store data source specific chi-square values

chi_sq_obs_by_source[s, d] <- sample_chi_obs_d

chi_sq_rep_by_source[s, d] <- sample_chi_rep_d

\# Add to overall totals

sample_chi_obs_total <- sample_chi_obs_total + sample_chi_obs_d

sample_chi_rep_total <- sample_chi_rep_total + sample_chi_rep_d

}

\# Store overall totals

chi_sq_obs_total[s] <- sample_chi_obs_total

chi_sq_rep_total[s] <- sample_chi_rep_total

}

\# Calculate overall Bayesian p-value

bayesian_pvalue_overall <- mean(chi_sq_rep_total >= chi_sq_obs_total)

\# Calculate data source specific p-values

bayesian_pvalues_by_source <- numeric(n_data)

for(d in 1:n_data) {

bayesian_pvalues_by_source[d] <- mean(chi_sq_rep_by_source[, d] >= chi_sq_obs_by_source[, d])

}

\# Create data source names if not available

source_names <- c("eDNA", "seine")

names(bayesian_pvalues_by_source) <- source_names

cat("Posterior predictive check complete!\n")

cat("Overall Bayesian p-value:", round(bayesian_pvalue_overall, 4), "\n")

cat("Data source specific p-values:\n")

for(d in 1:n_data) {

cat(" ", source_names[d], ":", round(bayesian_pvalues_by_source[d], 4), "\n")

}

\# Return comprehensive results

results <- list(

\# Overall results

bayesian.pvalue = bayesian_pvalue_overall,

chi_sq_obs_total = chi_sq_obs_total,

chi_sq_rep_total = chi_sq_rep_total,

mean_chi_sq_obs = mean(chi_sq_obs_total),

mean_chi_sq_rep = mean(chi_sq_rep_total),

\# Data source specific results

bayesian.pvalues.by.source = bayesian_pvalues_by_source,

chi_sq_obs_by_source = chi_sq_obs_by_source,

chi_sq_rep_by_source = chi_sq_rep_by_source,

mean_chi_sq_obs_by_source = apply(chi_sq_obs_by_source, 2, mean),

mean_chi_sq_rep_by_source = apply(chi_sq_rep_by_source, 2, mean),

\# Model info

n_samples = n_samples,

n_data_sources = n_data,

fit.stat = fit.stat,

source_names = source_names

)

return(results)

}
